# Thermal boldness: Volunteer exploration of extreme temperatures in *Drosophila melanogaster*

**DOI:** 10.1101/2021.10.15.464500

**Authors:** Carlos A. Navas, Gustavo A. Agudelo-Cantero, Volker Loeschcke

**Affiliations:** Department of Physiology, Institute of Biosciences, University of São Paulo. Rua do Matão 101, Tv 14, 05508-090 São Paulo, Brazil; Department of Biology - Genetics, Ecology and Evolution, Faculty of Natural Sciences, Aarhus University. Ny Munkegade 116, 8000 Aarhus C, Denmark

**Keywords:** Behavior, Critical Temperatures, *Drosophila*, Orientation, Thermal Biology

## Abstract

A dominant perception is that small and motile ectothermic animals must use behavior to avoid exposure to critical or sub-critical temperatures impairing physiological performance. Concomitantly, volunteer exploration of extreme environments by some individuals may promote physiological adjustments and enhance ecological opportunity. Here we introduce to the literature a Thermal Decision System (TDS) which is fully modular, thermally stable, versatile, and adaptable to study navigation through thermal landscapes in insects and other small motile animals. We used a specific setting of the TDS to investigate volunteer navigation through critical cold and hot temperatures in *Drosophila melanogaster*. We demonstrate that a thermally bold behavior (volunteer crossings through a Critical Temperature Zone, CTZ) characterized a fraction of flies in a sample, and that such a fraction was higher in an outbred population relative to isofemale lines. As set, the TDS generated a thermal gradient within the cold and hot CTZs, and the exploration of this gradient by flies did not relate simply with a tendency to be thermally bold. Mild fasting affected thermal exploration and boldness in complex manners, but thermal boldness was evident in both fasted and fed flies. Also, thermal boldness was not associated with individual critical temperatures. Finally, some flies showed consistent thermal boldness, as flies that performed an extreme thermal cross were more likely to perform a second cross compared with untested flies. We hypothesize that a simple “avoidance principle” is not the only behavioral drive for *D. melanogaster* facing extreme temperatures over space, and that this pattern may characterize other small motile ectothermic animals with analogous natural history. The physiological correlates, genetic architecture, and interspecific variation of thermal boldness deserve further consideration.

## 1. Introduction

The study of thermal adaptation in ectothermic animals incorporates principles from physiology, behavior, and ecology (Cowles and Bogert, 1944; Dawson and Templeton, 1963; Messenger, 1959). One early and broad-spectrum principle is the thermal dependence of many biological functions, an important factor in evolution. Thus, one approach to the study of thermal adaptation includes analyzing the relationship between biological functions and body temperature (thermal performance curves, TPCs), and the processes under which these curves evolve. In a typical case, a TPC depicts the thermal sensitivity of a given fitness-related trait over a range of body temperature and zero-performance values define the critical temperatures for activity (Huey and Slatkin, 1976). Then, the role of behavior in interpreting these curves is paramount because body temperature may constrain behavior and in turn, behavior has potential to affect or even modulate body temperature (Angilletta et al., 2006). Accordingly, the body temperature of ectothermic animals relates in a complex manner with environmental temperature given the interconnected influences of morphology, physiology, and behavior. For motile forms, and in manners conditioned by morphology and physiology, body temperature would be affected by patterns of navigation across thermal landscapes (Sears et al., 2016). Thus, if animals are granted self-producing (i.e., autopoietic) qualities and a cognitive domain (Thompson, 2007), their motility would include cognitive navigation through a thermal environment, and their orientation would follow decision rules to limit, avoid or even promote exposure to environmentally induced shifts in body temperature. These cognitive processes would also outline subtleties regarding thermal niche, a term used rather loosely in the biological literature (Gvoždík, 2018).

Orientation in thermal landscapes is influenced by the dynamics of temperature distribution across space and time (Sears et al., 2016), by social aspects of behavior including potential for aggregation (Aubernon et al., 2019) or heterospecific interactions (Winterová and Gvoždík, 2018), and by the feedback of behavioral decisions on physiology (Hutchison and Maness, 1979). This complexity highlights the general role of behavior as a driver for evolutionary shifts (Mayr, 1959, 1963), and the specific considerations of this postulate matter in the context of thermal adaption (Huey et al., 2003). However, incorporating the multifaceted nature of behavior into studies of thermal adaptation poses many challenges, and those linked to critical temperatures seem, so far, neglected. By definition, critical temperatures (minimum or *CT_min_*, and maximum or *CT_max_*) set thermal limits for field activity, and exceeding these temperatures can lead to behavioral impairment and lethality (Cowles and Bogert, 1944; Gunderson and Leal, 2016). A derived rationale, then, is that avoiding critical temperatures would be adaptive, as exposure to critical temperatures implies thermal risks (Andrew et al., 2013; Sunday et al., 2014). This is a compelling, well supported, and widely accepted postulate. Furthermore, the avoidance of critical temperatures is not only pervasive in motile animals but is perhaps the most ancestral type of thermoregulatory behavior (Nelson et al., 1984). However, Hutchison and Manes (1979) bring to debate a postulate of utmost importance: thermal landscapes, when explored behaviorally, create a dynamic physio-behavioral frame in which physiological adjustments such as hardening become possible. Therefore, navigation through thermal landscapes is both cause and consequence of individual thermal physiology.

It is possible that a simple “avoidance principle” may not be the only driver of evolution regarding navigation around critical temperatures in thermal landscapes. Even if the individual risk of approaching lethal temperatures seems ultimate, gains could exist in terms of cumulative hardening and enhanced ecological opportunity (e.g., short-term survival, shifting thermal niche, expanding animal distribution, finding resources, etc.) for some species (Hoffmann et al., 2007; Lee and Denlinger, 2010). Therefore, thermally risky behaviors may persist in a population, even if only at low densities. Such a possibility can occur in small ectothermic animals like many insects, whose motile forms can explore thermal landscapes, equilibrate rapidly with environmental temperatures, and reproduce in large numbers. With these considerations in mind, we set two primary goals with this study. First, we designed and tested a system to study exploration of extreme thermal landscapes in small insects and other motile ectothermic animals, which we introduce to the literature and describe in detail. Second, we take advantage of the versatility of the system to explore whether thermally risky behaviors, defined as the tendency to voluntarily enter and cross zones of critical temperatures (a behavior hereafter termed “thermal boldness”), exists in fruit flies of the species *Drosophila melanogaster*. We chose *Drosophila* because they meet the natural history criteria stated above, are experimentally versatile, and constitute a traditional model. Besides, previous findings encourage research on this topic. *Drosophila* lineages respond differently to experimental thermal regimes (Loeschcke et al., 1999), and in *D. melanogaster* laboratory fluctuating environments designed to mimic nature fail to reproduce wild adaptive patterns (Alton et al., 2017; Kellermann et al., 2015). This may be so because, compared to natural counterparts, experimental thermal environments depress spatial variance and reduce the relevance of navigation rules relative to other traits of thermal biology.

The scientific goal of this project is to obtain empirical evidence confirming that thermal boldness is a possible behavioral trait in small and motile ectothermic animals, using *D. melanogaster* as a model. At this point we do not engage into any hypothetic-deductive examination on the evolution of thermal boldness, a development we would consider premature for a foundational study. Specifically, we ask three related questions: i) Do flies voluntarily expose themselves to cold and hot critical temperatures? ii) If yes, does mild fasting enhance such thermally risky behaviors? and iii) Do fly lineages of different genetic makeup differ regarding inclination to approach critical temperatures? We anticipated that, if present, thermal boldness would characterize only a fraction of flies in a sample, and that such a fraction could be enhanced by mild fasting. Also, we supposed that potential differences in thermal boldness among fly lineages would validate future studies on its underlying genetic bases. Additionally, we explored –with no *a priori* hypotheses– a possible coupling between thermal physiology (temperature tolerance) and thermal boldness, and the possible consistency of this type of behavior for given individuals. Finally, we considered the possibility that, given the inequality of TPCs (Martin and Huey, 2008) and the less lethal nature of low critical temperatures, navigation towards or into extreme cold temperatures would be more common relative to navigation around or across extreme hot temperatures. We found that thermal boldness did characterize a fraction of individuals in both an outbred population and isofemale lines of *D. melanogaster*, with variation, nuances and unexpected patterns that are discussed in detail.

## 2. Material and Methods

### 2.1. Description of the system

#### 2.1.1. Overall design

The apparatus is depicted in Fig. 1, A-C and references to the parts cited in bold (Fig. 1A). Detailed measures of the system are provided in Fig. S1. Overall, the system has six compartments called “**Thermal Decision Systems**”, abbreviated “TDSs”, one of which was used to collect temperature data. The basic parts of each TDS are: 1) a **Home Bottle** at the base (which may or may not contain food), being a typical transparent stock bottle for *Drosophila* maintenance; 2) an assembly hereafter called **T-System** (Fig. 1A) given its “T” shape. The T-System connects to the home bottle through an ascending tube, and on top split into two symmetrical tubular structures of the same material and light diameter. Once the TDS was thermally stable, a 3) **Thermal Gradient** was conformed along the horizontal part of the T-System, and we use that term (or just “gradient”) to name this zone between the onset of CTZs. The thermal gradient is marked by the two black-rings placed 3.5 cm in both sides of the T-System (Fig. 1A). 4) Two **Temperature Coils**, located at each branch of the horizontal part of the T-System and externally delimited by black rubber rings, were responsible for creating both the thermal gradient and 5) the **Critical Temperature Zones** (CTZs), where cold and hot temperatures were most extreme (Fig. 1C). After the CTZs, and connected to each distal extreme of the horizontal part of the T-System, we placed 6) the **Feeding Bottles**, of the same type than the home bottle but always containing food. Finally, 7) a **Removable Stopper**, a tiny foam circle, was installed to temporally bar access from the home bottle to the T-System. A nylon thread was attached to the stopper and left the home bottle via a tiny perforation (Fig. 1A). This configuration served the purpose of easily allowing fly access by simply pulling the nylon thread, while causing virtually no disturbance to flies during removal (Video S1). The connections of the T-System to home and feeding bottles relied on perforated foam stoppers of the type used to lid *Drosophila* stock bottles (Fig. S1).

**Figure 1.**
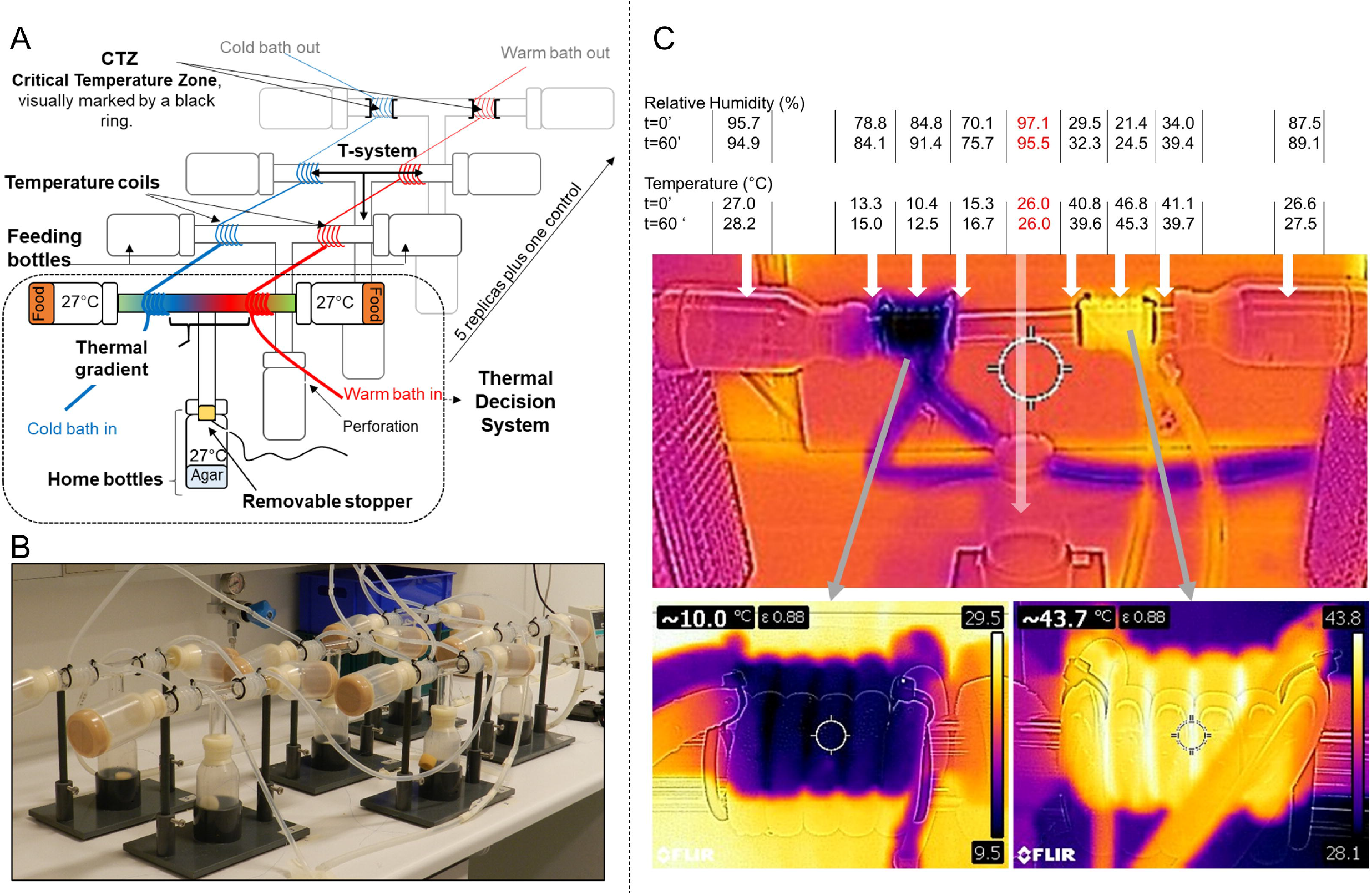
A) Schematic drawing of the system, showing 4 out of 6 replicates termed ‘Thermal Decision Systems’ (TDS), one of which served as control; B) Photograph of the system as installed; C) Thermographic images of an isolated TDS (upper panel) and each temperature coil (lower panel). The data shows average values of temperature and relative humidity for some key spots on the thermal gradient conformed in the upper part of the TDS. t=0’ refers to ‘time zero’, first event of data collection, and t=60’ refers to conditions one hour after. The numbers in red are estimates based in the temperature of the home bottle.

#### 2.1.2. Temperature control

Cooling and heating were provided simultaneously by using two independent laboratory thermal baths (NESLAB, RTE-300D for cooling; HETO, CBN 8-30 and HMT 200RS for heating) fit with a water-circulation system, creating the average thermal landscape depicted in Figure 1C. A silicon hose of 0.7 mm of internal diameter was fit to the outlet vent of each bath and then connected to the first TDS, as follows: each TDS was fit with a silicon coil of five loops at each side of the horizontal part of the T-System (Temperature Coils), placed 3.5 cm after the opening of the ascending tube, next to the black-ring working as a visual indicator (Fig. 1A, B, additional details in Fig. S1). The silicon coils connected TDSs in sequence via plastic tubes (1 cm). This set-up conferred both independent temperature control of cold and hot sides in each TDS and minimal temperature differences between TDS (conditions granted by a vast number of preliminary assays, see *Pilot studies* in Supplementary Material). Five TDSs were used simultaneously and one, always the front most as to reduce interference with observations, was fit with both temperature data loggers and a TC-08 Thermocouple Data Logger (Pico Technology) as to have online temperature records of the inner part of the tube surrounded by temperature coils. The central part of the portion of the inner tube bordered by the silicon hoses displayed the peak thermal barriers (see data table in Fig. 1C). Calibration tests allowed to estimate the inner temperature of the T-System under the influence of the silicone coils, with stable values at the target temperatures (Fig. S2). The system also created a non-planned gradient of relative humidity (Fig. 1C).

### 2.2. Flies maintenance and handling

Unless otherwise indicated, the data here reported refers to the fly lineage called Odd2010, which was derived from a wild population of *D. melanogaster* collected in October 2010 at Odder, South to Aarhus, Denmark (https://goo.gl/maps/yEG54oASEU5CynMB7). This lineage has been kept at 19°C since initial collection, with a generational time of 18 days. Additional tests were performed with isofemale lines of *D. melanogaster* from the Bloomington Drosophila Stock Center (ID number and genotype), which we chose randomly among those available in the laboratory as examples of highly inbred lines: 28240, **ISO40**, DGRP-812/RAL-812; 55014, **ISO14**, DRGP-31/RAL-31; 28213, **ISO13**, DRGP-589/RAL-589 (MacKay et al., 2012), hereafter abbreviated as lines 13, 14 and 40. All flies used were 3-4 days old, except when specifically noted. Flies were kept in 8 oz. (ca. 236.6 mL) stock bottles with 70 mL standard medium (5000 mL water, 200 g sugar, 150 g oatmeal, 80 g agar and 300 g yeast) at 25°C.

For each fly lineage, samples were composed by newly emerged flies of the same generation (i.e., breeding bottle). Briefly, we counted a low number of parents (ca. 30 pairs per breeding bottle) and applied routine transfers as to maintain comparable density among breeding bottles. For final fly selection, we first transferred flies from the breeding bottle to a fresh stock bottle (with food) using a systematic procedure leading to a presumably similar, yet uncounted, number of flies. From this new stock bottle, we aspirated about 50 flies using a silicone tubing with a glass point and a filter at one end, and then transferred them into a 7 mL empty vial (transfer vial) for later transfer. Then, we allowed 30 of these so aspirated flies to move upwards from the transfer vial to a home bottle to be installed in a TDS according to protocols. Flies to be fasted were transferred to home bottles with agar but no food 10 h before the onset of behavioral observations, always at daytime, whereas non-fasted flies were transferred to home bottles with food 60-90 min before data collection.

To achieve the final count of 30 flies passing from the transfer vial to home bottles, we used a small funnel fit to a silicon hose split by an aquarium air-valve that could be closed after the target fly number was attained. Thus, we controlled the number of flies for each sample, but not sex ratios. We opted for such a counting procedure because pilots showed that flies sorted under CO_2_ anesthesia 3-4 days after emergence moved less in the TDS than non-anesthetized flies (see *Preliminary tests* in Supplementary Material). Once the target fly number was attained, home bottles were transferred randomly to the TDSs. Occasionally up to two flies escaped or were squashed by the stopper in the setting of home bottles to T-Systems. These differences were ignored.

### 2.3. Behavioral observations and data collection

#### 2.3.1. Overall approach

The study was carried out at the fly lab of the Department of Biology - Genetics, Ecology and Evolution, Aarhus University. We turned the thermal baths on 70 min before the onset of data collection to let target temperatures stabilize in the system (see Supplementary Material for further details). Our overall thermal landscape (Fig. 1C) involved maintaining home bottles and ascending tubes at a room temperature of 26±1°C throughout the duration of the test. We started the test by removing the stoppers of all TDSs (< 5 sec) and granting flies access to T-Systems (Video S1). Flies reaching the horizontal part of T-Systems found an average thermal gradient ranging from ca. 15°C to 41°C, and CTZs with most extreme temperatures measured at ca. 10°C and 47°C. These values were stablished empirically and correspond to the most extreme temperatures at which we reported attempted crosses in preliminary tests (Fig. S2).

We observed fly behavior every 10 min and scored the behavioral variables described in the next section, all of them associated to activity and tendency to approach or enter the CTZs. Thus, 10 min after stoppers were removed, we collected behavioral data the first time (i.e., at time zero or *t_0_*) and kept collecting data at 10-min intervals. Given that the CTZs were extreme and could act as thermal barriers, we did not maintain a constant thermal setting. Rather, after ending a data collection cycle every 10 min, we increased (cold side) or decreased (hot side) settings by 0.5°C, so that the thermal configuration of the system slowly advanced from critical to subcritical. The option for a progressive change in temperature (as opposed to a fix setting) is part of the exploratory nature of this study. We anticipated that a temperature progression would lead to higher exploration and crossings for evaluation, and to enhanced analytical options (for example, regarding the temperatures at which a given fraction of the tested flies would cross). However, it turned out that most crosses were performed early in tests, as we discuss in the *Results*.

Cycles of data collection and thermal control were repeated up to A) two hours, *or* until B) at least 5% of flies (2 flies) in a sample had made a cold-cross *and* at least another 5% (2 flies) had made a warm-cross, *or* until C) at least one cross per side had occurred in each one of the TDSs (it turned out that only one test out of 8 was limited by time). Under protocol B, data collection on a given side, either cold or hot, was terminated when target crosses were reached. Then, we fixed the temperature at this first crossing side and maintained the temperature shift protocol only at the counterpart, until reaching target crosses or time. Cold and warm sides of TDSs were alternated among days in terms of left or right, relative to the system longitudinal axis, as placed on the working bench.

#### 2.3.2. Behavioral variables

At *t_0_*, and at 10-min intervals hereafter, we scored the number of flies i) exploring (i.e., moving) the COLD side of the T-system before the inner black-ring (*COLDEXP*); ii) exploring the WARM side of the T-system before the inner black-ring (*WARMEXP*); iii) touching the inner black-ring of the COLD side or inside the COLD-coiled area (*COLDCONTACT*); iv) touching the inner black-ring of the WARM side or inside the WARM-coiled area (*WARMCONTACT*); v) in the COLD-coiled area in atypical position and not moving (*CBI*); vi) in WARM coiled area in atypical position and not moving (*HEATCOM*); vii) in the feeding bottle after COLD coils (*COLDCROSS*); viii) in the feeding bottle after WARM coils (*WARMCROSS). CBI* stands for Cold-Induced Behavioral Impairment, a set of behavioral responses we observed in some flies attempting to cold-cross and that were reverted when temperature in the cold CTZ increase during our changing-temperature protocol (see *Results*). We avoid the term “chill coma” for it may suggest a physiological collapse to some readers, and this was not supported by further observations. We observed very few cases of *HEATCOM*, probably because of how flies crossed the hot CTZ (see *Results* on *Behavior after hot CTZ*), therefore, we did not analyze these variables formally but report anecdotally. Also, we observed few back-crossings from any feeding bottle to the thermal gradient (e.g., Video S2), but could not operationalize or analyze these occasional events.

#### 2.3.3. Consistency of behavior

To determine whether flies crossing CTZs could be generally more prone to thermal boldness as an individual trait, we performed a second test (next day) using cold-crossing or hot-crossing flies, according to a previous and first test. Here the protocol was modified slightly. First, we obtained crossing flies using 6 samples × 40 flies each, and set the CTZs at 44°C and 14°C (cold temperature elevated relative to original design as to avoid Cold-induced Behavioral Impairment or *CBI*, see *Behavior* in Results). With this thermal landscape configuration we obtained 40 Hot-crossing flies and 40 Cold-crossing flies (given the need of previous test, these flies were 4 days old). A third group of 40 non-previously tested flies (3 days old) was used as control. This procedure was repeated twice, so that we obtained 2 samples × 40 flies for each treatment (First Cold-crossing, First Hot-crossing, and Control). For final analyses we used total values (the sum of both tests) and compared the three groups of flies so treated in a test performed under identical conditions.

#### 2.3.4. Assumptions

The protocol here reported was based on these assumptions: i) despite the many stimuli that may coexist in the system, the number of flies leaving home bottles to circulate by (or even remain stationary at) any side of the T-System (cold or hot) relates to their behavioral inclination to explore the thermal gradient; ii) the number of flies inside CTZs or in feeding bottles after a CTZ cross relates to their tendency to explore critical or subcritical temperatures (cold or hot), i.e., is an indicator of thermal boldness; iii) eventual divergence in thermal exploration and boldness between fasted and not-fasted flies would result from enhanced motivation in fasted flies for exploring and crossing thermal barriers to get food (fasting enhanced activity in pilot tests, see *Preliminary tests* in Supplementary Material); iv) departure from symmetry in cold-crossers and hot-crossers indicates different inclination to explore cold and hot critical temperatures. In addition, some inferential statements in the Discussion assume that v) the navigation rules used by flies in the system somehow relate to those leading thermal exploration in nature. Also, we suppose that, as preliminary insights, vi) differences across lineages (e.g., outbred population vs. isofemale lines) in the inclination to navigate critical temperatures suggest a genetic basis for thermal boldness; and vii) consistent inter-individual variation in exploration of critical temperatures within lineages has basis on individual traits (e.g., genetic makeup, thermal history).

### 2.4. Measure of thermal tolerances

To explore possible associations between the tendency to perform an extreme temperature cross and physiological thermal tolerances, we planned a specific test based on 6 samples × 20 flies each (disregarding sex). Each of these six samples was associated to previous behavior in the testing system, as to have two samples of 20 hot-crossers, 20 cold-crossers and 20 non-crossers (i.e., flies remaining in the T-System at the end of a test). Flies composing these samples were obtained from behavioral experiments over two days, so that these thermal tolerances were measured on five-day old flies. We used one sample per treatment to test for the critical thermal minimum (*CT_min_*), the other for the critical thermal maximum (*CT_max_*). For testing, individual flies were distributed in small glass vials tightly sealed with plastic caps. We tested 60 flies (20 for each treatment) at once for a given critical temperature by immersing vials in a glass-made water bath (aquarium) allowing a clear view of each glass. To measure *CT_max_*, onset water temperature was 20°C, and then water temperature increased at a rate of 0.1°C/min. To measure *CT_min_*, onset bath temperature and rates of temperature change were the same, but the system contained a mixture of ethylene glycol and water in equal parts to avoid eventual freezing. Four observers collaborated with this measure by reporting end-temperatures for each fly according to typical behavioral observations, consisting mainly of flies falling down to the vial and showing no movement after tapping the vials gently with a metal stick. In that moment a fly was considered in thermal coma, and the temperature at the time was reported as the respective critical temperature.

### 2.5. Dry body mass

We measured dry body mass of individual flies, male and female. The procedure used follows Schou et al. (2015) and consisted in drying flies at 60°C for 24 h and then flash frozen them for later weighting. Flies were split in groups according to their behavior and feeding condition, and then stored in small containers with silica gel, as to avoid water absorption. Individual weight was measured with a Sartorius Laboratory balance (type MC5, Göttingen, Germany).

### 2.6. Data analysis

Although we report several variables and methods reaching some complexity, the research here reported is essentially inferential. We basically report keen observations of fly behavior and propose a biological hypothesis to explain them. We refer to statistical hypotheses when asking whether data display patterns according to standard statistical procedures. When pertinent we described temporal patterns of fly behavior along experiments, but this is exceptional and for most formal analyses the final number of flies exhibiting a given behavior is sufficient for proper inference. We applied parametric or non-parametric tests according to the assumption-wise profile of data sets. Briefly, we used either *t*-tests or Mann-Whitney U tests for comparing behavioral variables between two groups (e.g., fasted vs. non-fasted flies). For analyzing behavioral consistency among first crossers (cold or hot) and non-previously tested flies, we used Repeated Measures ANOVA, in this case to account for the temporal pattern of crossings within a given TDSs. To explore whether fly lineages differ in exploratory or bold behaviors we used General Linear Models (GLMs) followed by Bonferroni Post-hoc Tests (*BPHTs*). Physiological (*CT_min_* and *CT_max_*) and morphological (dry body mass) correlates of fly behavior among groups were tested via either GLMs or *t*-tests when applicable. All statistical analyses were performed in SPSS v. 22. In the main body of the paper, we provide type of analysis and level of significance, but placed full statistical details in the Supplementary Material, Table S1.

## 3. Results

### 3.1. Behavior

#### 3.1.1. Observation on fly behavior in the system

Our preliminary tests included numerous behavioral observations at room temperature that were performed with uncontrolled flies (for sex, density, and age), and are not suitable for a formal analysis. However, such observations were important to define our final procedure, and we report main conclusions as Supplementary Material. Briefly, flies tested at room temperature (no thermal gradient active) displayed diverse behaviors, with active flies that readily moved into the feeding bottles, more passive counterparts remaining in the home bottles, and many possible intermediate options. In formal tests with a thermal gradient, flies retained similar behaviors in the sense that some typically left the home bottle at the onset of the test, ascended the vertical tube of the T-System, and found the thermal gradient. Once in it, some flies remained stationary, other explored mainly one side of the gradient, and several explored the full gradient up to its limits, i.e., up to the onset of both CTZs. A fraction of the tested flies voluntarily approached CTZs and attempted either cold- or hot-crosses, and several flies died or were impaired in these attempts (Fig. S3).

Flies approached the hot CTZ by walking, advanced into it, but almost immediately made a sharp U-turn, escaping back into the gradient. Most flies barely surpassed the hot CTZ, but few entered it about 1 cm. At *t_0_*, i.e., at the highest temperatures, flies that did not U-turn after about 5 mm often switched to flight, sometimes hot-crossing erratically (Video S2), so that virtually all early hot-crosses occurred through this behavior. After *t_0_*, temperatures were slightly less extreme and flies increased the depth of initial advances into the hot CTZ, eventually reaching about 2 cm before a U-turn, or just not performing a U-turn at all and making a very rapid hot-cross by walking. Although rare observations, very few flies fell in heat coma (*HEATCOM*), few were found dead at the hot side and one fly died while attempting a hot-cross.

When approaching the cold CTZ, flies were exposed to about 15°C and theoretically retained physiological ability to U-turn, but only few flies displayed that behavior. Some approached the cold CTZ by slow walking and progressed into the cold CTZ exposing themselves to progressively lower temperatures. Other flies jut stopped by the cold CTZ boundary. Some flies that entered into the cold CTZ adopted atypical positions such as curved bodies, wings opened and legs upwards (Fig. S3), responses we referred to as Cold-induced Behavioral Impairment (*CBI*). Normally at *t_0_* several flies had attempted a cold-cross already, and at this time some flies under *CBI* accumulated within the cold CTZ (Fig.2, Fig. S3). However, most flies recovered from this condition as we increased temperatures in the cold side (Fig. 2). Finally, we observed two unmistakable cases of back crosses from the feeding bottle to the T-system through the cold CTZ.

**Figure 2.**
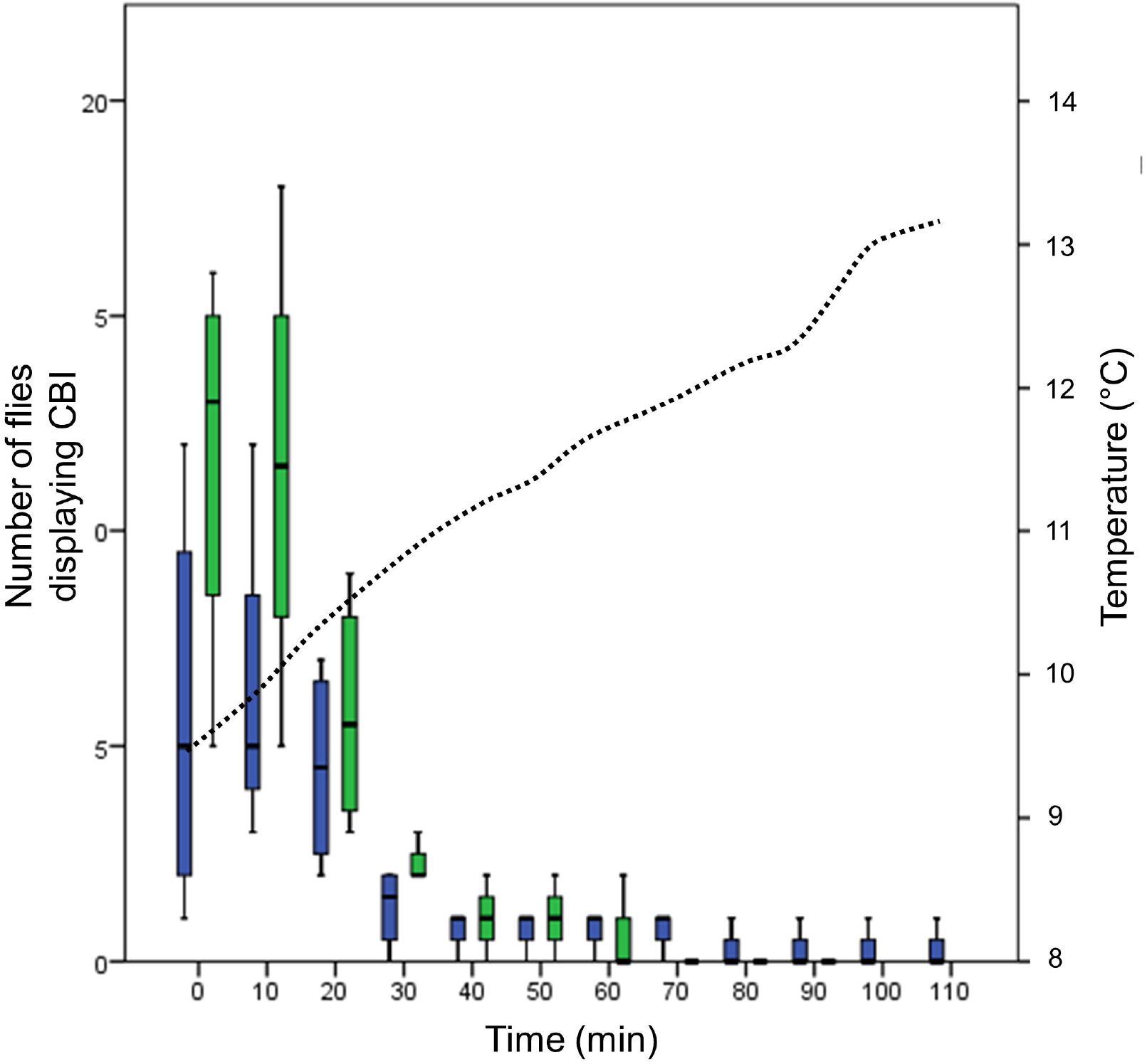
Average number of flies in Cold-induced Behavioral Impairment (*CBI*) as a function of time and temperature (right axis, dotted curve during tests. Values are non-cumulative over time. Green boxes for fasted (F) flies and blue boxes for non-fasted flies (NF). Boxes depict the first, second (median) and third quartiles containing 50% of data, and whiskers show the maximum and minimum values, except outliers (≥ 1.5 × IQR [the inter-quartile range] from the median) and extremes (≥ 3 × IQR from the median).

#### 3.1.2. Exploration of the thermal gradient

Regarding side selection in the gradient (cold vs. hot) by active fasted (F) or non-fasted (NF) flies, F flies stayed more often at the cold side of the gradient (*U-MW, P* < 0.01), while NF flies seemed to explore both sides similarly (*U-MW, P* = 0.051; *COLDEXP*, Fig. S4A; *WARMEXP*, Fig. S4B). Pooling all flies, active or stationary, the pattern was repeated and more flies stayed at the cold side of the gradient, independently of fasting condition (*t* test, *P* < 0.01). Apparently, the active exploration of the gradient did not display obvious patterns between fasting conditions, at least at the beginning of the experiment (Fig. S4A; Fig. S4B).

Regarding actual approaches to CTZs, F flies were bolder than NF flies when approaching the cold CTZ (*COLDCONTACT; U-MW, P* = 0.029; Fig S4C). However, this pattern lasted up to minute 10 (*t_10_*) and then diluted (Fig. S4C), partially because cold-crosses occurred or were attempted and led to *CBI* (Fig. 2). Then, the number of flies in condition to cross decreased with experimental time (Fig. 3). Fasted flies also approached the hot CTZ more often than NF flies throughout the test (*WARMCONTACT; U-MW, P* < 0.01; Fig. S4D). Overall, only a fraction of approaches to CTZs translated into successful crosses. Most flies exploring the thermal gradient approached CTZs, sometimes insistently, but did not attempt crosses.

**Figure 3.**
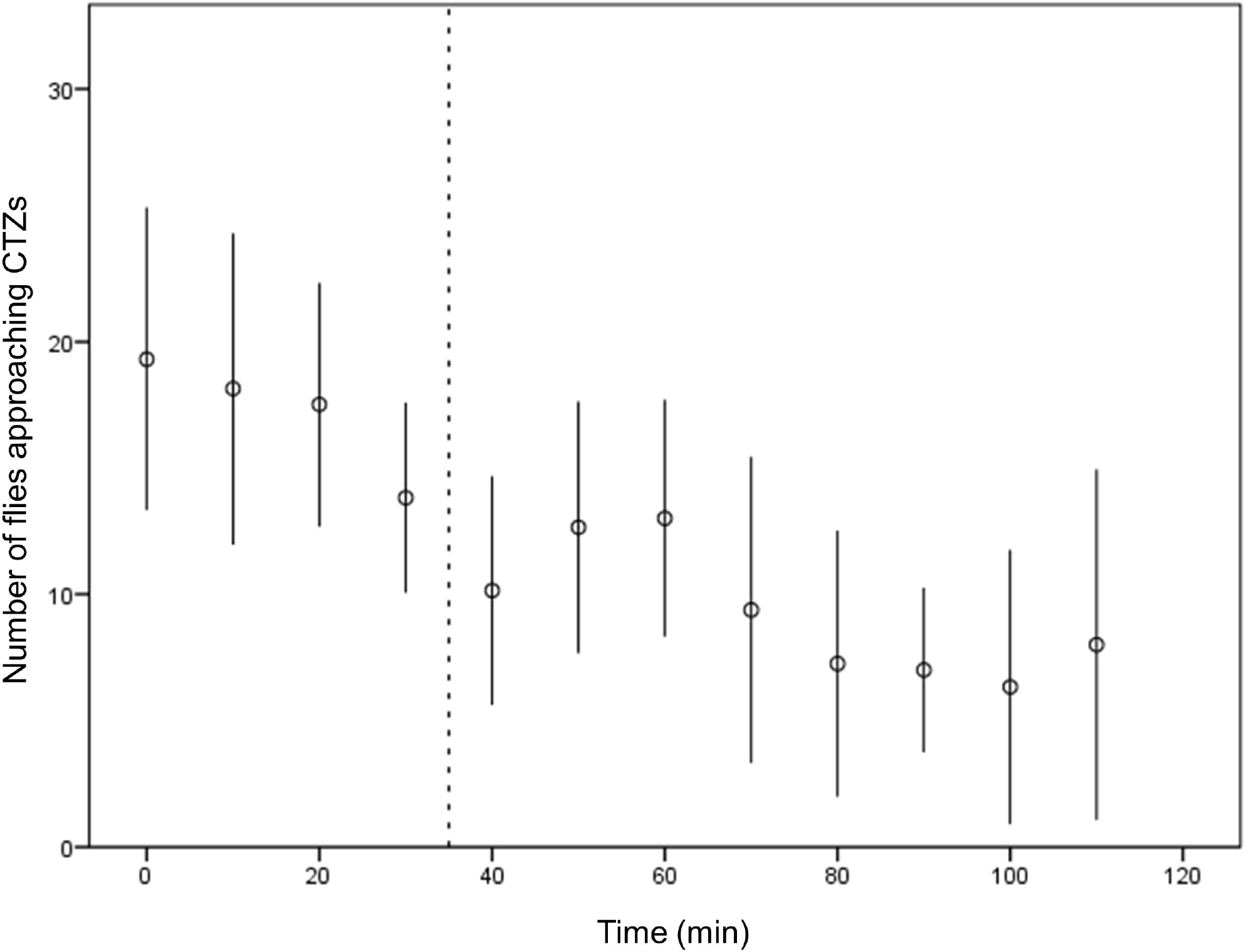
Number of flies approaching the onset of Critical Temperature Zones (CTZs, marked by black rubber rings in the T-System), i.e., summing up cold (*COLDCONTACT*) and hot (*WARMCONTACT*) approaches. Given a progressive reduction in exploratory drive and number of flies (due to both successful crosses and *CBI*), the total number of approaches declined after about 30 min (vertical dashed line).

#### 3.1.3. CTZ crosses

The mean time to meet protocol option B (5% crosses out of 30 flies in samples, see *Overall approach* in Methods) in cold crosses was shorter than the hot-cross equivalent, regardless fasting condition (cold-crosses, NF flies = 10.5 ± 1.82 min, F flies = 8.8 ± 0.85 min; hot-crosses, NF flies = 45.5 ± 1.35 min, F flies = 47.2 ± 1.32 min; see Table S2 for further details). In terms of cold-crosses, F flies displayed similar values than NF flies despite the former were bolder at exploring critical cold temperatures early in the experiment (*COLDCROSS; U-MW, P* = 0.966; Fig. 4A). A partial correlate for this pattern was that F flies displayed more cases of *CBI*, mostly early in the experiment (*U-MW, P* = 0.025; Fig. 2). On the other hand, more F flies appeared to cross through the hot CTZ relative to NF flies, but with considerable higher variation, so that no formal difference could be reported (*WARMCROSS; U-MW, P* = 0.833; Fig. 4B). Given that these patterns were heavily influenced by what happened up to *t_10_*, for an additional perspective we compared the isolated 10-60 min period of the experiment. During this time, F flies did show higher tendency to perform hot-crosses than NF flies (*U-MW, P* = 0.037), but the number of cold-crosses remained comparable among fasting groups (Fig. 4B).

**Figure 4.**
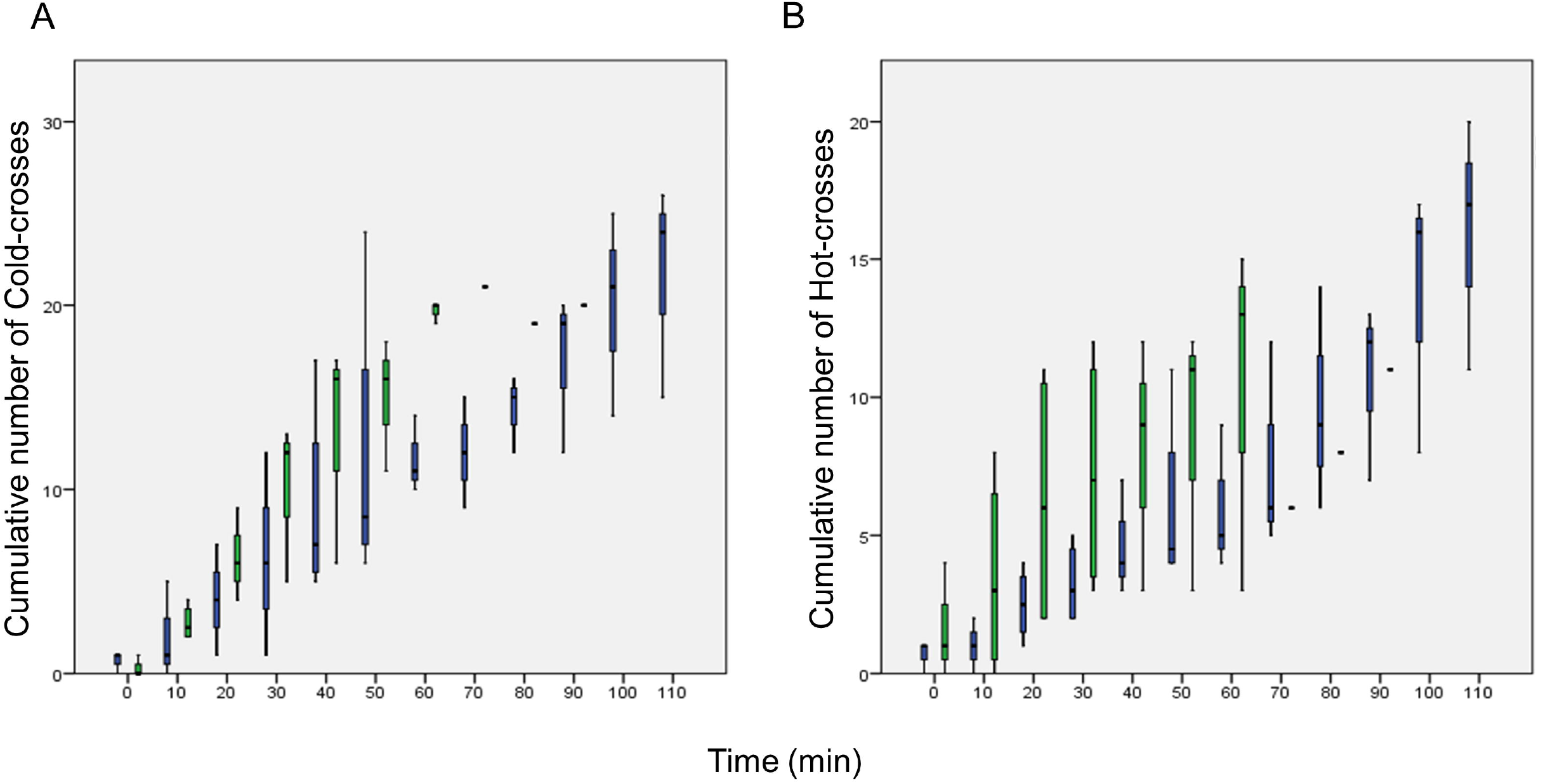
Cumulative number of crosses through extreme thermal barriers as a function of time. A) Successful cold-crosses, assessed by the number of flies in feeding bottles after the cold coils (*COLDCROSS*). B) Successful hot-crosses, assessed by the number of flies in feeding bottles after the warm coils (*WARMCROSS*). Figure details as in Fig. 2.

#### 3.1.4. Consistency of behavior

Sample-wise (we did not track individuals), flies that had performed a first extreme cross, cold or hot, displayed a tendency to accumulate more second crosses along time, relative to flies tested by their first time (Fig. 5A-B). The progression of second cold-crosses with time was clearly elevated in flies that had performed a first cold-cross relative to first hot-crossers and control flies (Repeated Measures ANOVA and *BPHT, P* < 0.01; Fig. 5A). A similar pattern was observed for originally hot-crossers performing a second hot-cross (Repeated Measures ANOVA, Crosses × Time, *P* = 0.045; Fig. 5B), but original cold-crossers also displayed a higher tendency to perform second hot-crosses relative to the control group (*BPHT, P* < 0.01).

**Figure 5.**
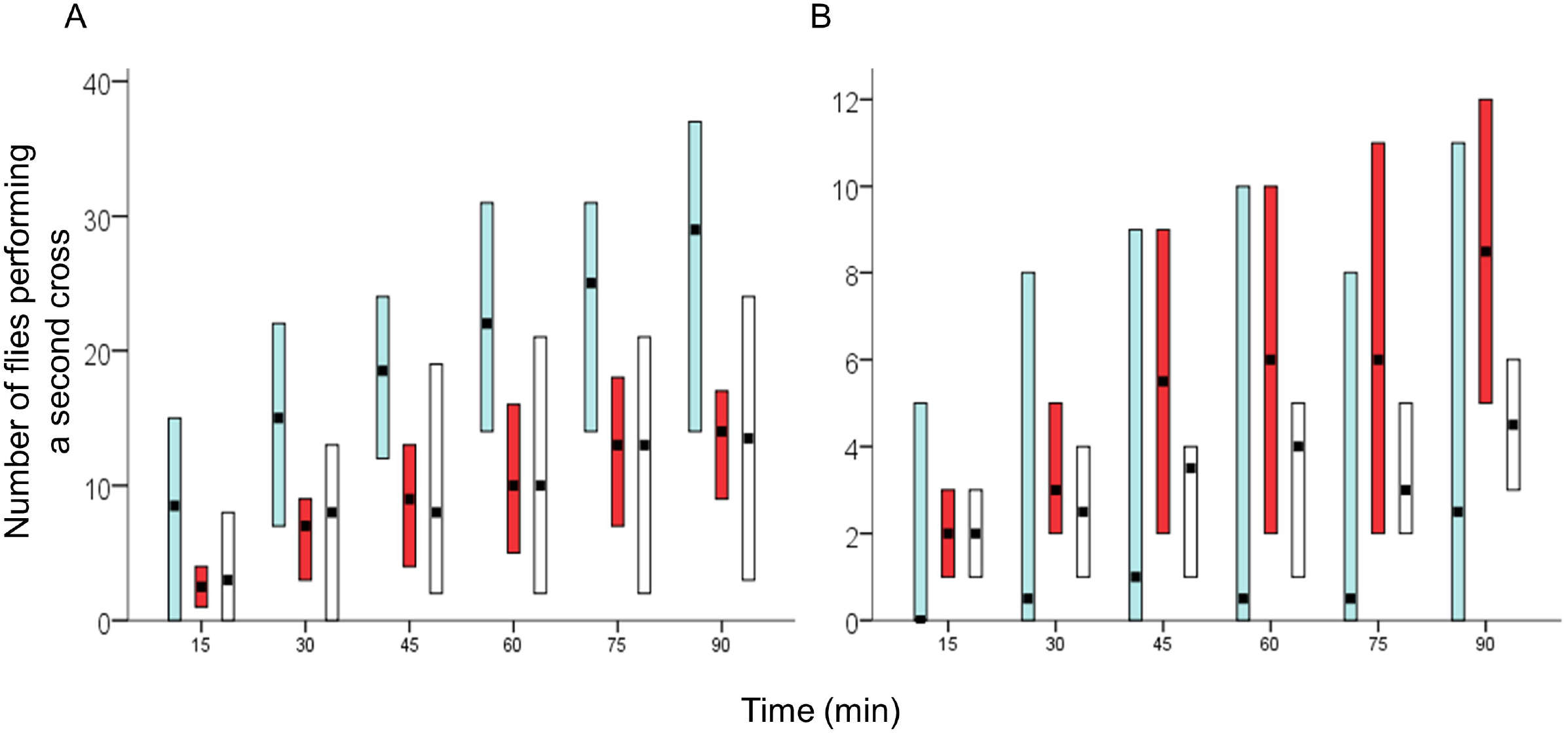
Count of flies performing a second cross through thermal barriers as a function of time, in a test for behavioral consistency (see main text for details). Bars show the minimum and maximum values, and the median out of four tests. A) shows the number of second cold-crosses performed by first cold-crossers (blue) or first hot-crossers (red), while B) displays comparable data for second hot-crosses. In both cases, second crosses are compared with a control group of non-previously tested flies (white), which by definition were tested only once for the purpose of these comparisons.

#### 3.1.5. Lineage-related differences

Overall, isofemale lines seemed behaviorally inhibited in terms of exploration of the thermal gradient, and their thermal boldness was low relative to that of Odd2010. For example, the number flies remaining in home bottles was higher for any isofemale line compared to Odd2010. Flies of the outbred population displayed more crosses through extreme thermal barriers (*TOTAL NUMBER OF CROSSES*; GLM, *P* < 0.01). Among isofemale lines, flies from line 14 performed more crosses through extreme thermal barriers compared to lines 13 and 40, which were more similar to each other (*BPHT, P* < 0.001; Fig. 6A, Fig. 6B). This pattern was mostly due to hot-crosses (Fig. 6D), particularly between 14 vs. 40 (*BPHT, P* = 0.041), whereas no differences among lines occurred for the very low values of cold-crosses (Fig. 6C). Despite pronounced differences in thermal boldness, flies of all lineages explored the cold CTZ similarly until the first 30 min of the test (*GLM, P* = 0.07), although line 13 exhibited less approaches to the cold CTZ relative to Odd2010 (*BPHT, P* < 0.01) but not to other lines (*BPHT, P* > 0.239). In contrast, Odd2010 flies approached the hot CTZ more often than any isofemale lines (*WARMCONTACT; BPHT, P* < 0.002 in all cases), but isofemale lines were also comparable (*BPHT, P* > 0.205).

**Figure 6.**
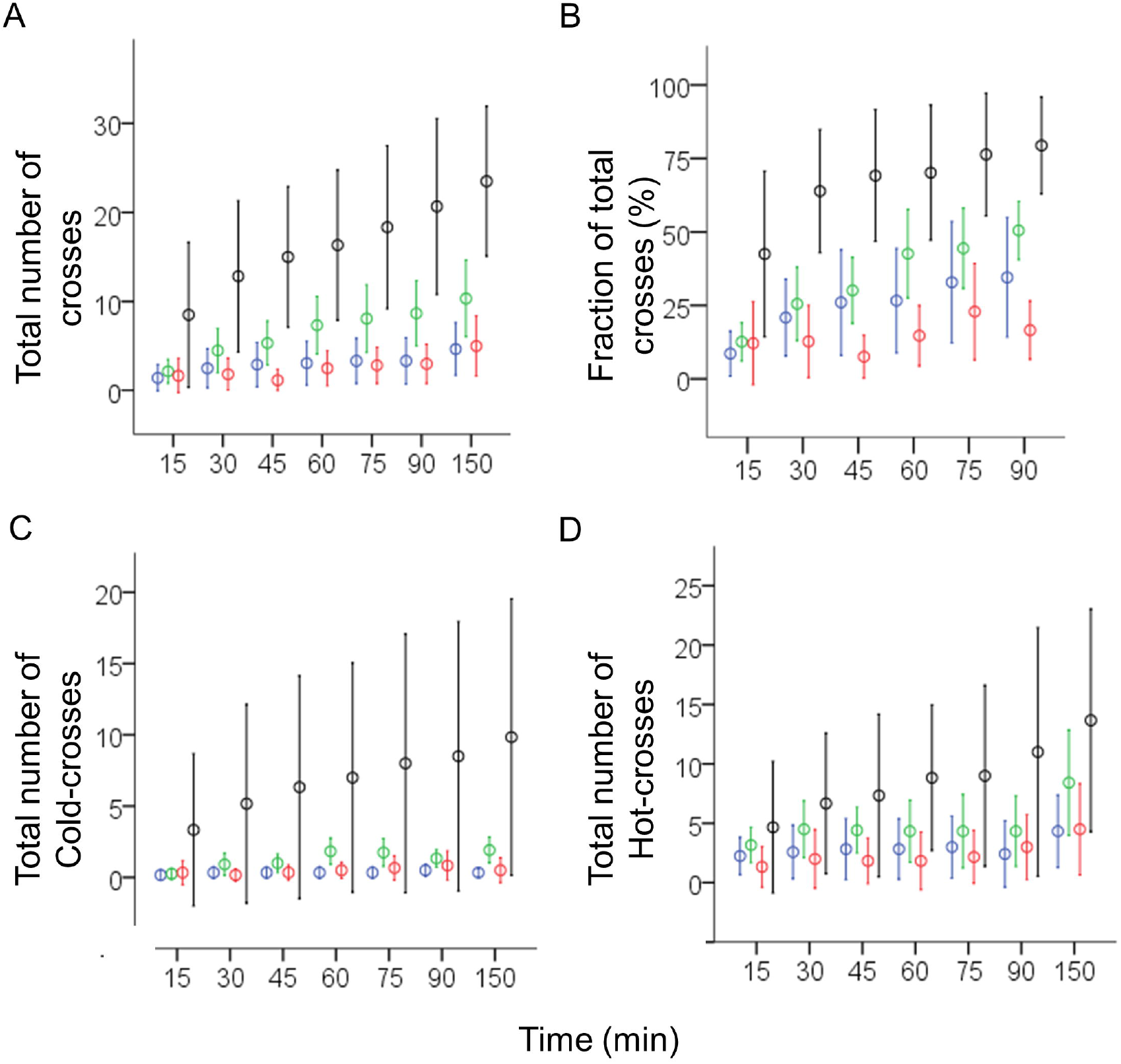
Lineage-related differences in crosses through extreme temperatures over time in *D. melanogaster*. The outbred Odd2010 population (black), core of most experiment, was compared with isofemale lines 14 (green), 13 (blue) and 40 (red) of the DGRP (see main text). A) Total number of extreme temperature crosses (i.e., *COLDCROSS* + *WARMCROSS*). B) fraction of total crosses in relation to the number of flies counted in the thermal gradient of the T-System. C) Total number of cold-crosses. D) Total number of hot-crosses. Circles and bars are the mean value ± 1 SD.

### 3.2. Thermal physiology and behavior

Cold-crossers were slightly less cold tolerant (i.e., higher *CT_min_*, measured only in Odd2010 flies after behavioral experiments) than other groups, yet not significantly (cold-crossers, *N* = 20, 6.6 ± 0.72°C; hot-crossers, *N* = 19, 6.15 ± 0.37°C; non-crossers, *N* = 19, 6.25 ± 0.56°C; GLM, *P* = 0.074). Regarding *CT_max_*, hot-crossers were less heat tolerant than cold-crossers, but similarly tolerant than non-crossers (hot-crossers, *N* = 20, 39.6 ± 0.75°C; cold-crossers, *N* = 20, 40.2 ± 0.36°C; non-crossers, *N* = 20, 39.9 ± 0.64°C; GLM, *P* = 0.011).

### 3.3. Morphological correlates

We measured dry body mass (*BM*, in mg) in a subsample of 352 flies, 195 females (139 NF and 56 F) and 157 males (112 NF and 45 F), collected after experiments. Full *BM* data appears in Table S3. As expected, females were on average 38% larger than males (females, *N*=195, *BM* = 0.274 ± 0.058 mg; males, *N*=157, *BM* = 0.198 ± 0.034 mg), and post-experimental F flies were about 13% smaller than NF flies (*BM-*F = 0.216 ± 0.045 mg; *BM*-NF = 0.248 ± 0.065 mg; *t*-test, *P* < 0.01). Cold-crossers and hot-crossers had comparable *BM* (cold-crossers, 0.256 ± 0.059 mg; hot-crossers, 0.245 ± 0.061 mg; *BPHT, P* = 0.207). Non-crosser flies were smaller than crosser flies (*BM*, 0.223 ± 0.061 mg; *BPHT, P* < 0.0001). A parallel pattern was found when comparing males only, but the trend was weaker among females.

## 4. Discussion

We present a modular, versatile and adaptable Thermal Decision System (TDS) to study the navigation of small and motile ectothermic animals through laboratory thermal landscapes. The system also allows an investigator to capture and isolate individuals that respond to a given set of decision rules, and so isolated individuals can be used for additional testing. Also, because the system is fully modular, researchers can choose the setup that is best suited to tackle a given research problem. For example, after making a given thermal cross, individuals could find another T-System with further settings, and so on. Finally, the contraption is not particularly expensive, can be set in a small climatic room, and has no special requirements. We provide all technical specifications as Supplementary Material, so that the system can be reproduced, modified, and enhanced.

Using one specific configuration of the system we confirmed that individual *D. melanogaster* voluntarily enter zones of critical temperatures, and that this is a populational phenomenon requiring the observation of many individuals. Our data show that 1) some flies voluntarily explore temperatures able to impair their behavior or even kill them; 2) this is not a common behavior and extreme temperatures act as barriers for most flies; 3) thermally bold individuals are more prone to engage in additional thermally risky behaviors; and 4) thermal boldness does not relate to thermal tolerance limits in any obvious manner. Because this is an introductory study, we refrain to incorporate thermal boldness into a theoretical evolutionary framework, a step that will develop as this behavior is further assessed. Also, we ignore how idiosyncratic this study is, for example given that both rearing and acclimation temperature may affect the thermal biology of *Drosophila* (Dillon et al., 2009).

Despite limitations, we infer that for small and motile ectothermic animals that reproduce in large numbers, a small fraction of individuals in a population might be thermally bold. If our results reflect intra-populational diversity (Wolf et al., 2007), as strongly supported by the outbred lineage, thermal boldness could typify the behavioral profile of a small fraction of individuals, along a continuum of risk-taking choices across thermal landscapes (Réale et al., 2007; Wilson et al., 1994, Fig. 7). Regarding underlying mechanisms, thermal boldness could have a genetic basis and be heritable, yet admitting contributions of the developmental environment, maternal effects, social influences, and other individual experiences (Falconer and Mackay, 1996). However, the behavioral differences among tested lineages of *D. melanogaster* favor the hypothesis of a genetic basis for thermal boldness. Although epigenetic sources of variation might be substrate for evolution (Burggren, 2016), differences in shyness and boldness in humans and other animals are mostly genetic and have proved heritable (Wilson et al., 1994 and citations therein), just as some decision-making behaviors of *D. melanogaster* like egg laying substrate selection (Miller et al., 2011). Finally, the notion of thermal boldness, as built upon our data, is compatible with examples of behavioral and physiological diversity within insect populations, including thermal biology, morphology (Forsman, 2000) and larval feeding behavior, as in the rovers vs. sitters *Drosophila* case (which has an autosomal basis, see Debelle and Sokolowski, 1987).

**Figure 7.**
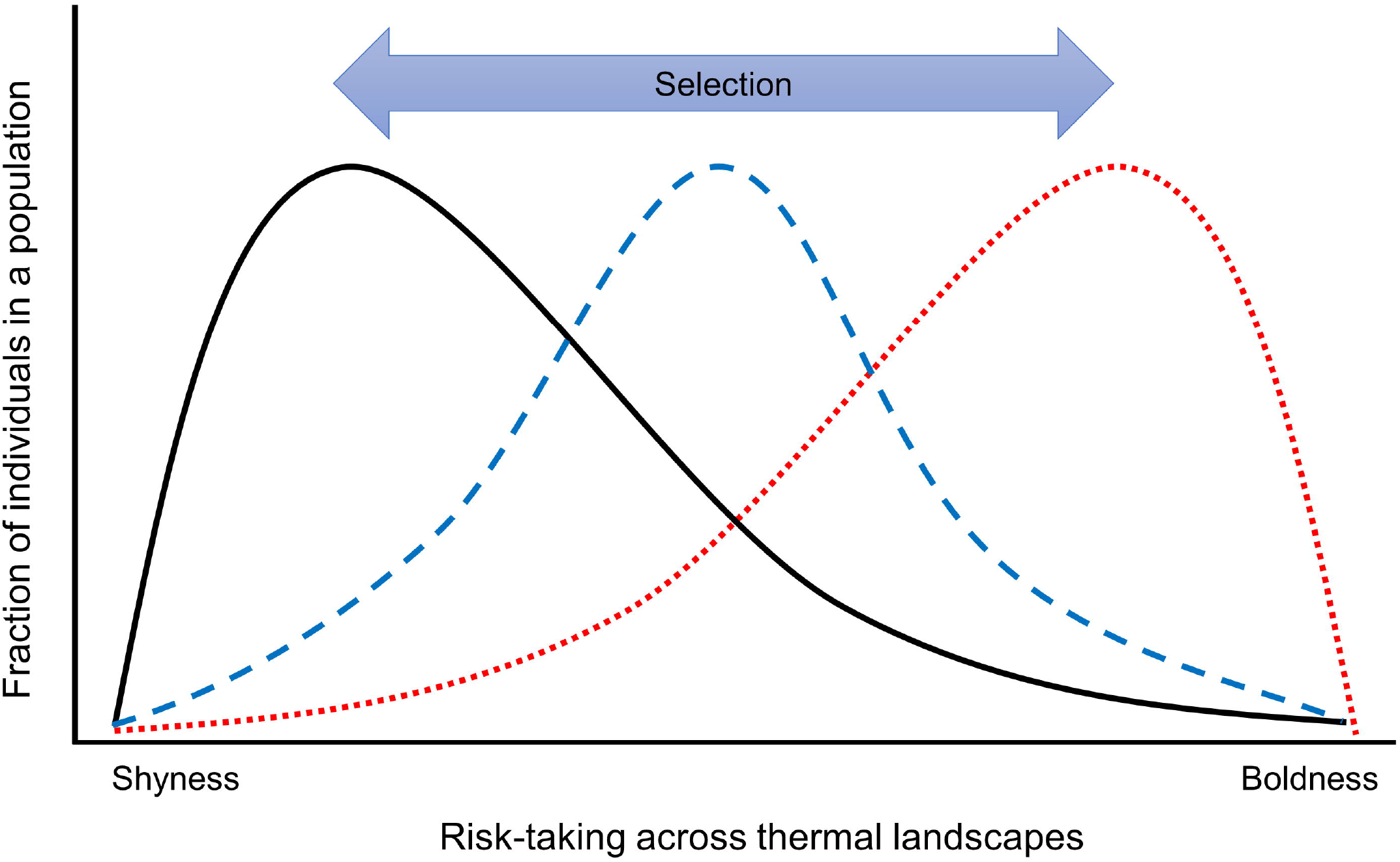
Hypothetic populational patterns of risk-taking behaviors across thermal landscapes in small motile ectothermic animals. Individuals exploring thermal landscapes may either retreat from extreme temperatures (shyness) or engage in their exploration (boldness). Polybehavioral populations could differ in the fraction of shy or bold individuals according to ecological and microevolutionary contexts. For instance, population dynamics at a given ancestral thermal niche (e.g., in range-core populations) may reflect directional selection favoring shy individuals (black solid line). Thermally bold individuals may be favored in novel environments with potential for colonization (e.g., edge populations, dotter red line). Transitional states would be possible in microevolutionary time.

In terms of ecological significance, thermal boldness could be linked to both ecological opportunity and impacts of thermal exposure on physiology (Hutchison and Maness, 1979; Terblanche et al., 2007), but the latter relationship remains to be established. Our data do not corroborate any obvious relationship between thermal boldness and physiological thermal tolerance, although admittedly this behavior could lead to cumulative hardening, perhaps realized in nature, given that some flies were persistent in their volunteer exposure to critical temperatures (e.g., Video S1). Albeit previous studies have suggested little relation between thermal tolerance and some behavioral traits of *D. melanogaster* (e.g., locomotor activity, feeding behavior and place memory) (Bahrndorff et al., 2016; Gioia and Zars, 2009), the relationship between thermal physiology and behavior may vary when thermal variation along space is involved (Salachan et al., 2021). However, experimental selection studies have prioritized thermal variations in time, but natural fluctuations involve both time, space, and navigational possibilities. Perhaps this fact explains why attempts to mimic natural thermal fluctuations have failed to replicate the adaptive trends observed in the wild (Kellermann et al., 2015).

The onset temperatures used in this study were based on empirical identification of the coldest and warmest temperatures at which crosses occurred (Supplementary Material). So, at least from this perspective, both CTZs were “similarly extreme”. Despite this care, exploration and thermal boldness were asymmetrical regarding cold and hot extremes, a pattern perhaps related to physiological risk. As set in our system, cold temperatures impaired but did not kill flies, contrary to hot temperatures (even if at low frequencies), a finding perhaps capturing one aspect of the asymmetrical nature of TPCs (Martin and Huey, 2008). Even so, fly behavior at and across the hot CTZ suggest that the upper thermal limits of TPCs, as typically measured, do not necessarily relate to the impossibility to explore such thermal zones (Fig. 8). Collectively, the experimental system presented in this paper provides new options to study the relationships between TPC structure and individual thermal physiology and behavior, with specific nuances for cold and hot extremes. Also, in the context here reported, flies immobilized within the cold CTZ did recover when temperature increased. These flies may have entered chill coma but only if understood as a reversible physiological state (Hazell and Bale, 2011), though our observations did not support that possibility.

**Figure 8.**
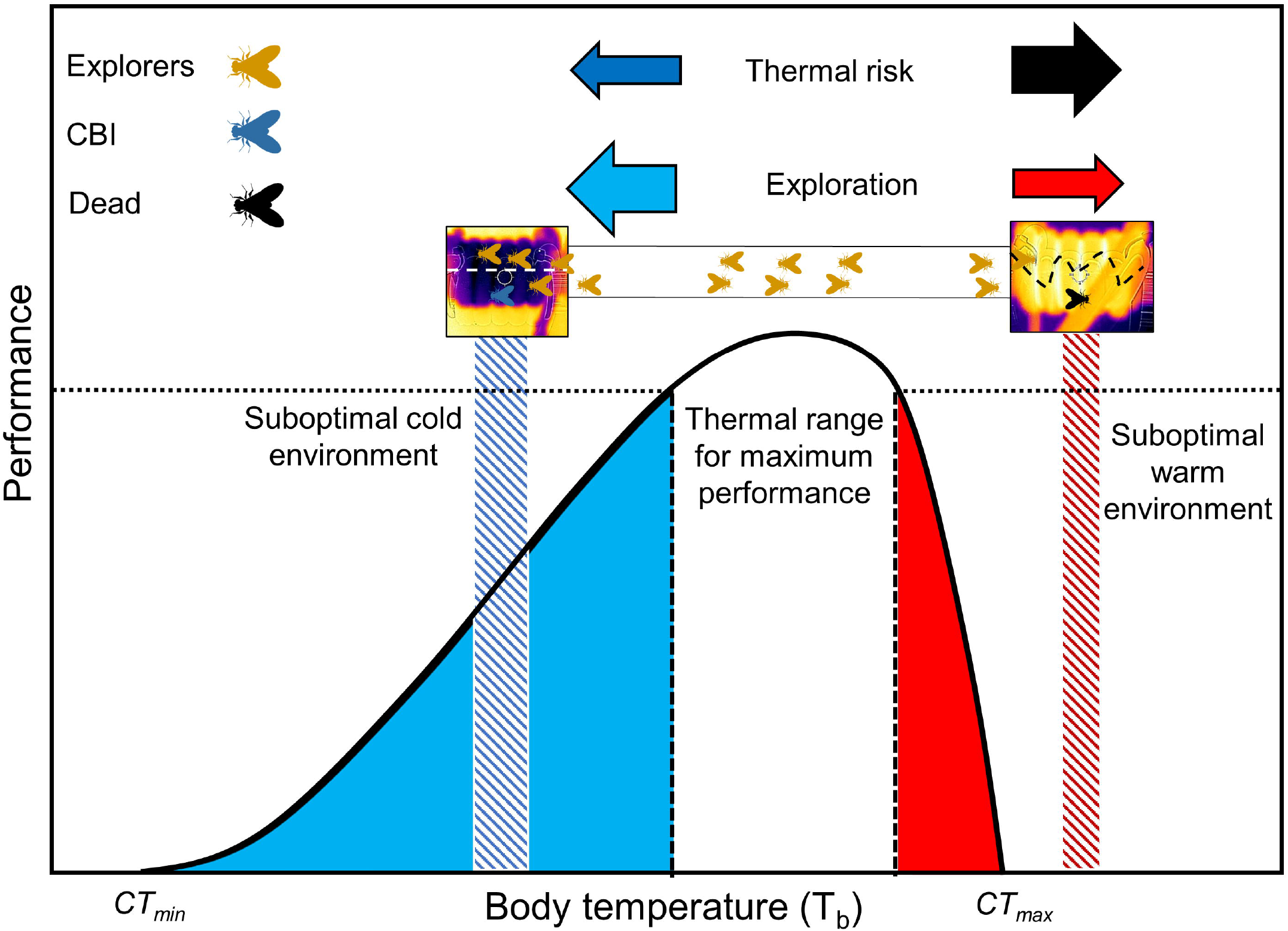
Hypothetical Thermal Performance Curve (TPC) highlighting cold-to-hot asymmetry (Martin and Huey, 2008). In this figure hatched areas represent thermal barriers based on present data (4-6°C above mean *CT_min_*, 5-6°C above mean *CT_max_*. Physiological risk would be asymmetrical (dark blue vs. black arrows), from temporarily impaired activity at the cold end (blue fly) to dying individuals at the hot end (black fly). This asymmetry may explain higher exploration of extreme cold by *D. melanogaster* (light blue vs. red arrows), as well as the contrast between voluntarily walking into inhibiting cold (dashed white line within cold barrier) and erratic flying at the hot side (dashed black line within heat barrier). However, *CT_min_* and *CT_max_*, as typically assessed, do not necessarily reflect limits for voluntary exploration and thermal boldness.

Finally, mild fasting enhanced exploration and thermal boldness, particularly at the hot side, with more pronounced effects soon after release into the TDS. Enhanced exploration and boldness were expected given that fasting may elicit more risky behaviors (Moran et al., 2021). In parallel, the behavior of first crossers could have affected other flies via odor clues because fasting also enhances odor-tracking behavior in *Drosophila* (Farhadian et al., 2012). Another caveat is that odor-tracking behaviors may be collaterally affected by temperature, for example through differential dynamics of odor clues at each side of the thermal gradient. Despite these uncertainties, thermal boldness was expressed at both cold and hot sides and was well-defined also in non-fasted flies, even if at lower frequencies. Thus, we propose that thermal boldness may be enhanced by some types of ecological risk, but it is not exclusive to such scenarios. In this sense, our results may not reflect proclivity to forage on substrates above critical temperatures, such as in the ant species *Iridomyrmex purpureus* (Andrew et al., 2013).

## 5. Conclusions

In small and motile ectothermic animals, thermal physiology relates in complex manners with orientation in thermal landscapes. This is evident in the flexibility and limits of TPCs (Angilletta et al., 2002; Navas, 2006), their response to experimental selection (Huey and Kingsolver, 1993), the diversity among measures of performance (Kellermann et al., 2019), and the impact of level of organization (Rezende and Bozinovic, 2019). These analyses assume optimality (Martin and Huey, 2008) and the generalization that critical temperatures, which by definition cause physiological and ecological damage, must be behaviorally avoided by individuals (Andrew et al., 2013; Sunday et al., 2014). Although this perception is supported by a strong theory, the avoidance of critical temperatures, at least as typically measured, may be less pervasive than originally thought. Alternative behaviors, including thermal boldness, may be perpetuated given potential links with physiological adjustment and ecological opportunity. Thermally bold individuals could pioneer the expansion of distribution into some new adaptive zones, as reported for some species during the first stages of invasion (Lindström et al., 2013; Mayr, 1963; Wright et al., 2010). Although it is clear that *D. melanogaster* exhibits thermal boldness, the generality of this behavior needs to be further scrutinized in other species, as well as its consistency, heritability, and evolutionary potential.

## Authorship contribution statement

**Carlos A. Navas:** Conceptualization, Methodology, Validation, Investigation, Formal analysis, Writing, review & editing, Visualization, Funding acquisition (travel). **Gustavo A. Agudelo-Cantero:** Writing, review & editing, Visualization. **Volker Loeschcke:** Conceptualization, Methodology, Writing, review & editing, Project administration, Funding acquisition.

## Acknowledgments

We thank John Svane Jensen, Assistant Engineer at the Department of Biology - Zoophysiology, Aarhus University, for his outstanding contribution to fabricate the system. We are grateful to Trine Bech Søgaard and Annemarie Højmark for technical help in the fly lab, and to Jesper Givskov Sørensen for general support and discussion. We thank the Danish Natural Sciences Research Council (FNU, grant 4002-00113B) for financial support to VL, the State of São Paulo Science Foundation, FAPESP, for financial support to CAN (FAPESP No. 2014/16320-7) and GAAC (FAPESP No. 2019/23325-9), and the Aarhus University Research Foundation for supporting the visit of CAN to Aarhus.

## References

Alton, L.A., Condon, C., White, C.R., Angilletta, M.J., 2017. Colder environments did not select for a faster metabolism during experimental evolution of Drosophila melanogaster. Evolution (N. Y). 71, 145–152. https://doi.org/10.1111/evo.13094

Andrew, N.R., Hart, R. a., Jung, M.P., Hemmings, Z., Terblanche, J.S., 2013. Can temperate insects take the heat? A case study of the physiological and behavioural responses in a common ant, Iridomyrmex purpureus (Formicidae), with potential climate change. J. Insect Physiol. 59, 870–880. https://doi.org/10.1016/j.jinsphys.2013.06.003

Angilletta, M.J., Bennett, A.F., Guderley, H., Navas, C.A., Seebacher, F., Wilson, R.S., 2006. Coadaptation: A unifying principle in evolutionary thermal biology. Physiol. Biochem. Zool. 79, 282–294. Doi 10.1086/499990

Angilletta, M.J., Niewiarowski, P.H., Navas, C.A., 2002. The evolution of thermal physiology in ectotherms. J. Therm. Biol. 27, 249–268. https://doi.org/10.1016/S0306-4565(01)00094-8

Aubernon, C., Hedouin, V., Charabidze, D., 2019. The maggot, the ethologist and the forensic entomologist: Sociality and thermoregulation in necrophagous larvae. J. Adv. Res. 16, 67–73. https://doi.org/10.1016/j.jare.2018.12.001

Bahrndorff, S., Gertsen, S., Pertoldi, C., Kristensen, T.N., 2016. Investigating thermal acclimation effects before and after a cold shock in Drosophila melanogaster using behavioural assays. Biol. J. Linn. Soc. 117, 241–251. https://doi.org/10.1111/bij.12659

Burggren, W., 2016. Epigenetic inheritance and its role in evolutionary biology: Re-evaluation and new perspectives. Biology (Basel). 5. https://doi.org/10.3390/biology5020024

Cowles, R.B., Bogert, C.M., 1944. A preliminary study of the thermal requirements of desert reptiles. Bull. Am. Museum Nat. Hist. 83, 261–296. https://doi.org/10.1086/394795

Dawson, W.R., Templeton, J.R., 1963. Physiological responses to temperature in the lizard, Crotaphytus coilaris. Physiol. Zool. 36, 219–236. https://doi.org/10.1086/physzool.36.3.30152308

Debelle, J.S., Sokolowski, M.B., 1987. Heredity of Rover Sitter: Alternative foraging strategies of Drosophila melanogaster larvae. Heredity (Edinb). 59, 73–83. DOI 10.1038/hdy.1987.98

Dillon, M.E., Wang, G., Garrity, P.A., Huey, R.B., 2009. Thermal preference in Drosophila. J. Therm. Biol. 34, 109–119. https://doi.org/10.1016/j.jtherbio.2008.11.007

Falconer, D.S., Mackay, T.F.C., 1996. Introduction to Quantitative Genetics, 4th ed, Trends in Genetics. Longman, Harlow, England.

Farhadian, S.F., Suarez-Farinas, M., Cho, C.E., Pellegrino, M., Vosshall, L.B., 2012. Post-fasting olfactory, transcriptional, and feeding responses in Drosophila. Physiol. Behav. 105, 544–553. https://doi.org/10.1016/j.physbeh.2011.09.007

Forsman, A., 2000. Some like it hot: Intra-population variation in behavioral thermoregulation in color-polymorphic pygmy grasshoppers. Evol. Ecol. 14, 25–38. Doi 10.1023/A:1011024320725

Gioia, A., Zars, T., 2009. Thermotolerance and place memory in adult Drosophila are independent of natural variation at the foraging locus. J. Comp. Physiol. a-Neuroethology Sens. Neural Behav. Physiol. 195, 777–782. https://doi.org/10.1007/s00359-009-0455-2

Gunderson, A.R., Leal, M., 2016. A conceptual framework for understanding thermal constraints on ectotherm activity with implications for predicting responses to global change. Ecol. Lett. 19, 111–120. https://doi.org/10.1111/ele.12552

Gvoždík, L., 2018. Just what is the thermal niche? Oikos. https://doi.org/10.1111/oik.05563

Hazell, S.P., Bale, J.S., 2011. Low temperature thresholds: Are chill coma and CTmin synonymous? J. Insect Physiol. 57, 1085–1089. https://doi.org/10.1016/j.jinsphys.2011.04.004

Hoffmann, A.A., Sørensen J.G., Loeschcke, V., 2003. Adaptation of Drosophila to temperature extremes: bringing together quantitative and molecular approaches. J. Therm. Biol. 28: 175–216. https://doi.org/10.1016/S0306-4565(02)00057-8

Huey, R.B., Hertz, P.E., Sinervo, B., 2003. Behavioral drive versus behavioral inertia in evolution: a null model approach. Am. Nat. 161, 357–366. https://doi.org/10.1086/346135

Huey, R.B., Kingsolver, J.K., 1993. Evolution of resistance to high temperature in ectotherms. Am. Nat. 142, S21–S46. https://doi.org/10.1086/285521

Huey, R.B., Slatkin, M., 1976. Cost and benefits of lizard thermoregulation. Q. Rev. Biol. 51, 363–384. https://doi.org/10.1086/409470

Hutchison, V.H., Maness, J.D., 1979. The role of behavior in temperature acclimation and tolerance in ectotherms. Am. Zool. 19, 367–384. https://doi.org/10.1093/icb/19.1.367

Kellermann, V., Chown, S.L., Schou, M.F., Aitkenhead, I., Janion-Scheepers, C., Clemson, A., Scott, M.T., Sgro, C.M., 2019. Comparing thermal performance curves across traits: how consistent are they? J. Exp. Biol. 222. https://doi.org/10.1242/jeb.193433

Kellermann, V., Hoffmann, A.A., Kristensen, T.N., Moghadam, N.N., Loeschcke, V., 2015. Experimental evolution under fluctuating thermal conditions does not reproduce patterns of adaptive clinal differentiation in Drosophila melanogaster. Am. Nat. 186, 582–593. https://doi.org/10.1086/683252

Lee, R.E., Denlinger, D.L., 2010. Rapid cold-hardening: ecological significance and underpinning mechanisms, in: Denlinger, D.L., Lee, R.E. (Eds.), Low Temperature Biology of Insects. Cambridge University Press, Cambridge, UK, pp. 35–58

Lindström, T., Brown, G.P., Sisson, S.A., Phillips, B.L., Shine, R., 2013. Rapid shifts in dispersal behavior on an expanding range edge. Proc. Natl. Acad. Sci. 110, 13452–13456. https://doi.org/10.1073/pnas.1303157110

Loeschcke, V., Bundgaard, J., Barker, J.S.F., 1999. Reaction norms across and genetic parameters at different temperatures for thorax and wing size traits in Drosophila aldrichi and D. buzzatii. J. Evol. Biol. 12, 605–623. https://doi.org/10.1046/j.1420-9101.1999.00060.x

MacKay, T.F.C., Richards, S., Stone, E.A., Barbadilla, A., Ayroles, J.F., Zhu, D., Casillas, S., Han, Y., Magwire, M.M., Cridland, J.M., Richardson, M.F., Anholt, R.R.H., Barrón, M., Bess, C., Blankenburg, K.P., Carbone, M.A., Castellano, D., Chaboub, L., Duncan, L., Harris, Z., Javaid, M., Jayaseelan, J.C., Jhangiani, S.N., Jordan, K.W., Lara, F., Lawrence, F., Lee, S.L., Librado, P., Linheiro, R.S., Lyman, R.F., MacKey, A.J., Munidasa, M., Muzny, D.M., Nazareth, L., Newsham, I., Perales, L., Pu, L.L., Qu, C., Ràmia, M., Reid, J.G., Rollmann, S.M., Rozas, J., Saada, N., Turlapati, L., Worley, K.C., Wu, Y.Q., Yamamoto, A., Zhu, Y., Bergman, C.M., Thornton, K.R., Mittelman, D., Gibbs, R.A., 2012. The Drosophila melanogaster Genetic Reference Panel. Nature 482, 173–178. https://doi.org/10.1038/nature10811

Martin, T.L., Huey, R.B., 2008. Why “Suboptimal” is optimal: Jensen’s inequality and ectotherm thermal preferences. Am. Nat. 171, E102–E118. https://doi.org/10.1086/527502

Mayr, E., 1963. Animal species and evolution. Harvard University Press, Cambridge, Massachussets.

Mayr, E., 1959. The emergence of evolutionary novelties, in: Tax, S. (Ed.), Evolution after Darwin. University of Chicago Press, Chicago.

Messenger, P.S., 1959. Bioclimatic studies with insects. Annu. Rev. Entomol. Palo Alto 4, 183–206. https://doi.org/10.1146/annurev.en.04.010159.001151

Miller, P.M., Saltz, J.B., Cochrane, V.A., Marcinkowski, C.M., Mobin, R., Turner, T.L., 2011. Natural Variation in Decision-Making Behavior in Drosophila melanogaster. PLoS One 6, e16436. https://doi.org/10.1371/journal.pone.0016436

Moran, N.P., Sánchez□Tójar, A., Schielzeth, H., Reinhold, K., 2021. Poor nutritional condition promotes high□risk behaviours: a systematic review and meta analysis. Biol. Rev. 96, 269–288. https://doi.org/10.1111/brv.12655

Navas, C.A., 2006. Patterns of distribution of anurans in high Andean tropical elevations: Insights from integrating biogeography and evolutionary physiology. Integr. Comp. Biol. 46, 82–91. https://doi.org/10.1093/icb/icj001

Nelson, D.O., Heath, J.E., Prosser, C.L., 1984. Evolution of temperature regulatory mechanisms. Am. Zool. 24, 791–807. https://doi.org/10.1093/icb/24.3.791

Réale, D., Reader, S.M., Sol, D., McDougall, P.T., Dingemanse, N.J., 2007. Integrating animal temperament within ecology and evolution. Biol. Rev. 82, 291–318. https://doi.org/10.1111/j.1469-185X.2007.00010.x

Rezende, E.L., Bozinovic, F., 2019. Thermal performance across levels of biological organization. Philos. Trans. R. Soc. B-Biological Sci. 374. https://doi.org/10.1098/rstb.2018.0549

Salachan, P. V, Sorensen, J.G., Maclean, H.J., 2021. What can physiological capacity and behavioural choice tell us about thermal adaptation? Biol. J. Linn. Soc. 132, 44–52. https://doi.org/10.1093/biolinnean/blaa155

Sears, M.W., Angilletta, M.J., Schuler, M.S., Borchert, J., Dilliplane, K.F., Stegman, M., Rusch, T.W., Mitchell, W.A., 2016. Configuration of the thermal landscape determines thermoregulatory performance of ectotherms. Proc. Natl. Acad. Sci. 113, 10595–10600. https://doi.org/10.1073/pnas.1604824113

Schou, M.F., Loeschcke, V., Kristensen, T.N., 2015. Inbreeding depression across a nutritional stress continuum. Heredity 115, 56–62. https://doi.org/10.1038/hdy.2015.16

Sunday, J.M., Bates, A.E., Kearney, M.R., Colwell, R.K., Dulvy, N.K., Longino, J.T., Huey, R.B., 2014. Thermal-safety margins and the necessity of thermoregulatory behavior across latitude and elevation. Proc. Natl. Acad. Sci. U. S. A. 111, 5610–5. https://doi.org/10.1073/pnas.1316145111

Thompson, E., 2007. Mind in Life: Biology, Phenomenology, and the Sciences of Mind. Harvard University Press.

Wilson, D.S., Clark, A.B., Coleman, K., Dearstyne, T., 1994. Shyness and boldness in humans and other animals. Trends Ecol. Evol. 9, 442–446. https://doi.org/10.1016/0169-5347(94)90134-1

Winterová, B., Gvoždík, L., 2018. Influence of interspecific competitors on behavioral thermoregulation: developmental or acute plasticity? Behav. Ecol. Sociobiol. 72, 169. https://doi.org/10.1007/s00265-018-2587-2

Wolf, M., van Doorn, G.S., Leimar, O., Weissing, F.J., 2007. Life-history trade-offs favour the evolution of animal personalities. Nature 447, 581–584. https://doi.org/10.1038/nature05835

Wright, T.F., Eberhard, J.R., Hobson, E.A., Avery, M.L., Russello, M.A., 2010. Behavioral flexibility and species invasions: the adaptive flexibility hypothesis. Ethol. Ecol. Evol. 22, 393–404. https://doi.org/10.1080/03949370.2010.505580

