## Supplementary Material for "Thermal boldness: Volunteer exploration of extreme temperatures in *Drosophila melanogaster*"

### Pilot studies

#### 1. Startup temperature

To gather information about what temperature to use as thermal barriers we run a pilot with an uncontrolled sample of flies bred from wild type. These flies were randomly collected from breeding stock with no control on age, sex, or any other biological parameter. We run preliminary tests with such fly samples ($N$ = 30) starting with thermal barriers set at 14.5 and 43.5°C respectively, with a 0.5°C change (higher at the warm side and lower at the cold side). This procedure was repeated until 61 TDS were tested in wild-type flies, group in which less than 5% would cross barriers set to 47.5°C and 9.5°C. (Fig. S2). Thus, we set initial temperatures at 47°C (hot side) and 9°C (cold side) after proving that these induced either death by overheating or behavioral impairment due to cold. This thermal gradient was produced by five loops of silicon tubing (type) placed in the same position among TDS (temperature coils, see Fig. 1 in the main text); the first cold or warm loop was placed exactly at 3.5 cm distance from the ascending tube (Fig. S1). This point was marked by black rubber rings serving the double purpose of fixing the silicon tubing and marking the steep gradient point. The portion of the temperature coils under direct influence of the first silicon loop, either cold or hot (CTZs, Fig. 1), produced a steep gradient of about 5°C across less than 0.5 cm.

#### 2. Preliminary tests

We run preliminary tests in flies less than 3-4 days old, varying the counting system of flies (collected after $CO_2$ anesthesia versus mouth pumping), their feeding history (ad libitum to 15h fasting), and the influence of different number of flies in home bottles. These tests were initially ran with thermal baths off, so that temperature across the system virtually followed room temperature, and let us know that: 1) $CO_2$-treated flies moved less in the TDS than untreated flies; 2) among many combinations of feeding treatments, maximum activity (yet "normal", as assessed by a human observer) was elicited at about 10 h of food deprivation; 3) flies required time to settle down after transfer from breeding stock to the home bottle of the TDS; and 4) in the absence of a thermal gradient (i.e., observational system at a room temperature ca. 26°C), some very active flies ended up easily in feeding bottles), whereas more passive flies remained in home bottles. Given these considerations, we opted for transferring flies to home bottles the day before of tests, so that on the test day no additional bottle-transfer was necessary. We used agar but not media in home bottles, as to produce mild fasting, with transferring protocols calibrated so that fasting time started the last night before tests in flies already inside home bottles (with stoppers activated), in ready-to-go TDSs. In this protocol we refer hereafter to **time zero ($t_0$)** as the time 10 min after the last stopper was removed, that is, the time at which the first observation was made once flies gained free access to the T-system. The thermal baths were turned at least 70 min before $t_0$, so that the system had plenty of time to equilibrate thermally. To start data collection, then, the only necessary step was to remove the stopper preventing fly access to the T-system (Video S1). This proved an excellent alternative because all removable stoppers could be removed in about 5 sec, granting an almost simultaneous start in the five operational TDSs. These preliminary tests required about two months of dedicated work. Preliminary tests with a temperature gradient *on* began only after, with the data reported in Figure S1.

### 3. **Gradient stability tests**

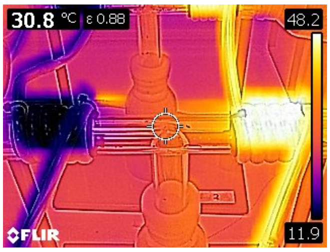


The stability of the system was tested via thermography (calibrated for proper emissivity) and applied to 6 independent tests performed on different days. Each test consisted of i) turning the gradient on, ii) let the system stabilize for 70 min (anticipating this proposed timing for real tests) and iii) capture thermographic images of each one of the TDS (see figure to the right as an example), iv) tabulate the external temperature (see below) at the core of each temperature coil and v) compare the distribution of values. The full calibration procedure included 48 thermographic images, and different set-points with mean cold-coil temperatures ranging from 6.5 °C to 11.3 °C and mean hot-coil temperatures ranging from 47.9°C to 51.0°C. At the onset of pilots, the maximum difference in temperature among TDSs were 2.3°C (cold) and 4.2 (hot), but adjustments mainly in water circulation speed, reduction of hose length and added isolation of bath exist lines with foam significantly reduced this variation. Several arrangements were tested, and by the end of calibration the four best arrangements (including the one used and illustrated in this paper) led to maximum differences of 1.0°C to 1.3°C (cold) and 0.8°C to 1.1°C (hot) among the TDSs, at bath nominal temperatures near those used in tests, with a range of standard deviations as follows: cold, 0.367°C to 0.564°C; hot, 0.337°C to 0.417°C. It is important to note that temperatures registered via thermography relate to external temperatures of the TSDs, i.e., on the immediate surface of target spots (e.g., silicone hoses), and that those external temperatures may differ from the actual temperatures experienced by flies inside the T-System. Therefore, part of the calibration included relating the nominal temperatures set in the thermal baths to internal temperatures in the middle of heating/cooling coils (Fig. S5), where thermal barriers occur (Fig. 1C, main text).

### SUPPLEMENTARY TABLES

**Table S1.** Statistical details on fly behavior, specifying the response variable and groups. For every behavioral variable, the number of flies was compared between or among groups. *U-MW* stands for the Mann-Whitney U test, RM is repeated measures for ANOVA, and GLM is a General Linear Model with a resulting *Z* statistic (according to the SPSS coding). The symbol "***" stands for a *P-value* < 0.0001. Comparisons up to time 90 min (unless otherwise stated), given too many empty cells in further readings.

| Comparison (variable, groups) | Test | Statistic | *P-value* |
|---|---|---|---|
| *Side selection* by active fasted flies, cold vs. hot | *U-MW* | 276.5 | *** |
| *Side selection* by active non-fasted flies, cold vs. hot | *U-MW* | 425.5 | 0.051 |
| *Side selection* by all fasted flies (active plus inactive), cold vs. hot | *t* | 4.068 | *** |
| *Side selection* by all non-fasted flies (active plus inactive), cold vs. hot | *t* | 5.609 | *** |
| *COLDCONTACT*, fasted vs. non-fasted flies | *U-MW* | 769.5 | 0.029 |
| *WARMCONTACT*, fasted vs. non-fasted flies | *U-MW* | 997.5 | *** |
| *COLDCROSS*, fasted vs. non-fasted flies | *U-MW* | 591.5 | 0.966 |
| *CBI*, fasted vs. non-fasted flies | *U-MW* | 771.5 | 0.025 |
| *WARMCROSS* (whole test), fasted vs. non-fasted flies | *U-MW* | 605.5 | 0.833 |
| *WARMCROSS* (10-60 min of test), fasted vs. non-fasted flies | *U-MW* | 329.5 | 0.037 |
| *SECOND COLDCROSS,* first cold-crossers vs. first hot-crossers vs. control (Crosses × Time) | RM ANOVA, $F_{10,45}$ | 2.894 | 0.007 |
| *SECOND WARMCROSS,* first hot-crossers vs. first cold-crossers vs. control (Crosses × Time) | RM ANOVA, $F_{10,45}$ | 2.094 | 0.045 |
| *TOTAL NUMBER OF CROSSES*, Odd2010 vs. isofemale lines | GLM, *Z* | 66.98 | *** |
| *COLDCROSS*, Odd2010 vs. isofemale lines | GLM, *Z* | 45.16 | *** |
| *WARMCROSS*, Odd2010 vs. isofemale lines | GLM, *Z* | 17.82 | *** |

| | | | |
|---|---|---|---|
| *COLDCONTACT*, Odd2010 vs. isofemale lines | GLM, *Z* | 6.228 | *** |
| *COLDCONTACT* (up to minute 30), Odd2010 vs. isofemale lines | GLM, *Z* | 2.463 | 0.07 |
| *WARMCONTACT*, Odd2010 vs. isofemale lines | GLM, *Z* | 11.101 | *** |
| $CT_{min}$, cold-crossers vs. hot-crossers vs. non-crossers | GLM, *Z* | 2.737 | 0.074 |
| $CT_{max}$, hot-crossers vs. cold-crossers vs. non-crossers | GLM, *Z* | 4.877 | 0.011 |
| *Dry mass*, fasted vs. non-fasted flies | *t* (unequal variances) | 4.765 | *** |
| *Dry mass*, males vs. females | *t* (unequal variances) | -15.43 | *** |
| *Dry mass*, hot-crossers vs. cold-crossers vs. non-crossers | GLM, *Z* | 9.521 | *** |

**Table S2**. Descriptive statistics comparing the time (in minutes) needed to terminate tests in Odd2010 flies, non-fasted (NF) and fasted (F). Because tests ended according to target crosses or time, these values indicate how prone flies were to engage in extreme thermal crossings. *N* is the number of tests, each one consisting of simultaneous testing in 5 TDSs. Minimum (*Min*), maximum (*Max*) and mean (*Mean*) time were averaged among the 5 TSDs for each test. Reported values are the average *Min*, *Max* and *Mean* for the four tests.

| **Group:** | | ***N*** | ***Min*** | ***Max*** | ***Mean*** | ***SD*** |
|---|---|---|---|---|---|---|
| NF | Full Test | 4 | 50.0 | 110.0 | 95.0 | 30.00 |
| | Cold-Cross | 4 | 8.5 | 12.5 | 10.5 | 1.82 |
| | Hot-Cross | 4 | 44.5 | 47.5 | 45.5 | 1.35 |
| F | Full Test | 4 | 30.0 | 90.0 | 60.0 | 24.49 |
| | Cold-Cross | 4 | 8.0 | 10.0 | 8.8 | 0.85 |
| | Hot-Cross | 4 | 45.5 | 48.5 | 47.2 | 1.32 |

**Table S3**. Dry body mass of a subsample of tested flies (total *N* = 352) across groups, expressed in mg.

| **Thermal Behavior** | **Feeding Condition** | **Sex** | ***N*** | ***Min*** | ***Max*** | ***Mean*** | ***SD*** |
|---|---|---|---|---|---|---|---|
| Cold-crossers | Non-fasted | F | 47 | 0.212 | 0.464 | 0.296 | 0.057 |
| | | M | 34 | 0.175 | 0.328 | 0.228 | 0.035 |
| | Fasted | F | 20 | 0.167 | 0.310 | 0.257 | 0.038 |
| | | M | 14 | 0.111 | 0.224 | 0.186 | 0.031 |
| Hot-crossers | Non-fasted | F | 42 | 0.209 | 0.428 | 0.295 | 0.052 |
| | | M | 28 | 0.156 | 0.262 | 0.190 | 0.031 |
| | Fasted | F | 20 | 0.173 | 0.299 | 0.240 | 0.033 |
| | | M | 11 | 0.154 | 0.256 | 0.198 | 0.027 |
| Non-crossers | Non-fasted | F | 50 | 0.165 | 0.485 | 0.272 | 0.065 |
| | | M | 50 | 0.130 | 0.258 | 0.193 | 0.028 |
| | Fasted | F | 16 | 0.129 | 0.299 | 0.225 | 0.044 |
| | | M | 20 | 0.124 | 0.195 | 0.174 | 0.020 |

**Supplementary Videos.**

**Video S1.** Stopper removal granted fly access from the home bottle to the T-System and marked the onset of behavioral observations. This step was conducted in less than 5 sec for all Thermal Decision Systems.
File name: THERMAL_BOLDNESS_SUPP_VIDEO_1.

**Video S2.** An adult fly of *D. melanogaster* (18 h of starvation at 26°C before the test) crossing voluntarily a hot thermal barrier (~41°C) four times. A first forward cross (*WARMCROSS*, from the thermal gradient to the feeding bottle) was already made by the time 0 sec when we started recording the video, but this fly opted for keep exploring the area between the outer black ring and the feeding bottle instead of entering directly to the last one. Then, at second 6 the fly entered the hot CTZ again and crossed backward to the thermal gradient (first *BACKWARM*) before second 9 was completed. After approximately 10 seconds of exploration, the fly approached again the hot CTZ, entered and crossed (second *WARMCROSS*). Two seconds after (second 26), the fly entered again the hot CTZ and backcrossed the second time, staying in the thermal gradient until the end of the video. All crosses performed by this fly, both forward and backward, occurred in less than 3 sec through an erratic flying activity. The underlying reasons behind this behavior and the decision of not entering the bottle containing food in the first place are unknown.
File name: THERMAL_BOLDNESS_SUPP_VIDEO_2.

### Captions for Supplementary Figures

**Figure S1**. Measurement details of components of the Thermal Decision System introduced in this article to study behavior and navigation of small mobile insects in extreme thermal landscapes.

**Figure S2**. Plot illustrating the fraction of flies, out of samples of 30 flies, crossing thermal barriers up to 15 min after $t_0$. This fraction is plotted as a function of the estimated temperature of thermal barriers, as for the data-logging in the control TDS.

**Figure S3**. Photographs illustrating a fraction of the T-System under the influence of the cold coils (cold CTZ), set at about 9°C in a pilot on adult flies of the Odd2010 lineage randomly obtained from breeding stock (uncontrolled sex or age). Flies approaching this cold end rarely attempted to fly and walked increasingly slower. Some flies attempting to cross fell into Cold-induced Behavioral Impairment (bottom right, see main text for more details). This information was used to set onset conditions in formal tests with controlled fly stock.

**Figure S4**. Number of flies exploring the thermal gradient before CTZs (*COLDEXP* or *WARMEXP*) or approaching CTZs (*COLDCONTACT* or *WARMCONTACT*) as a function of time during tests. Active flies exploring either the cold (A) or the warm (B) side of the gradient. Flies actually approaching the cold CTZ (C) or the hot CTZ (D). Green boxes for fasted (F) flies and blue boxes for non-fasted flies (NF). Boxes depict the first, second (median) and third quartiles containing 50% of data, and whiskers show the maximum and minimum values, except outliers (≥ 1.5 × IQR [the inter-quartile range] from the median) and extremes (≥ 3 × IQR from the median).

**Figure S5**. Calibration curve for the system. The Y axis is temperature of data-loggers inside the tube of the T-System, in the middle of the area under influence of the heating/cooling coils (i.e., the peak of thermal barriers). The X axis presents thermal bath nominal temperature. Temperatures at the thermal barriers differed about 2°C from nominal bath temperatures and correlation was high ($P <$ 0.001 and $R^2 > 0.8$ in both cases). The equations relating the predicted temperatures at thermal barriers ($Y$) for each side of the T-System, assuming a linear fit, were: $WARM\colon Y = 0.83X + 6.194$ and $COLD\colon Y = 0.897X + 3.189$, where $X$ is the respective nominal bath temperature in each case.

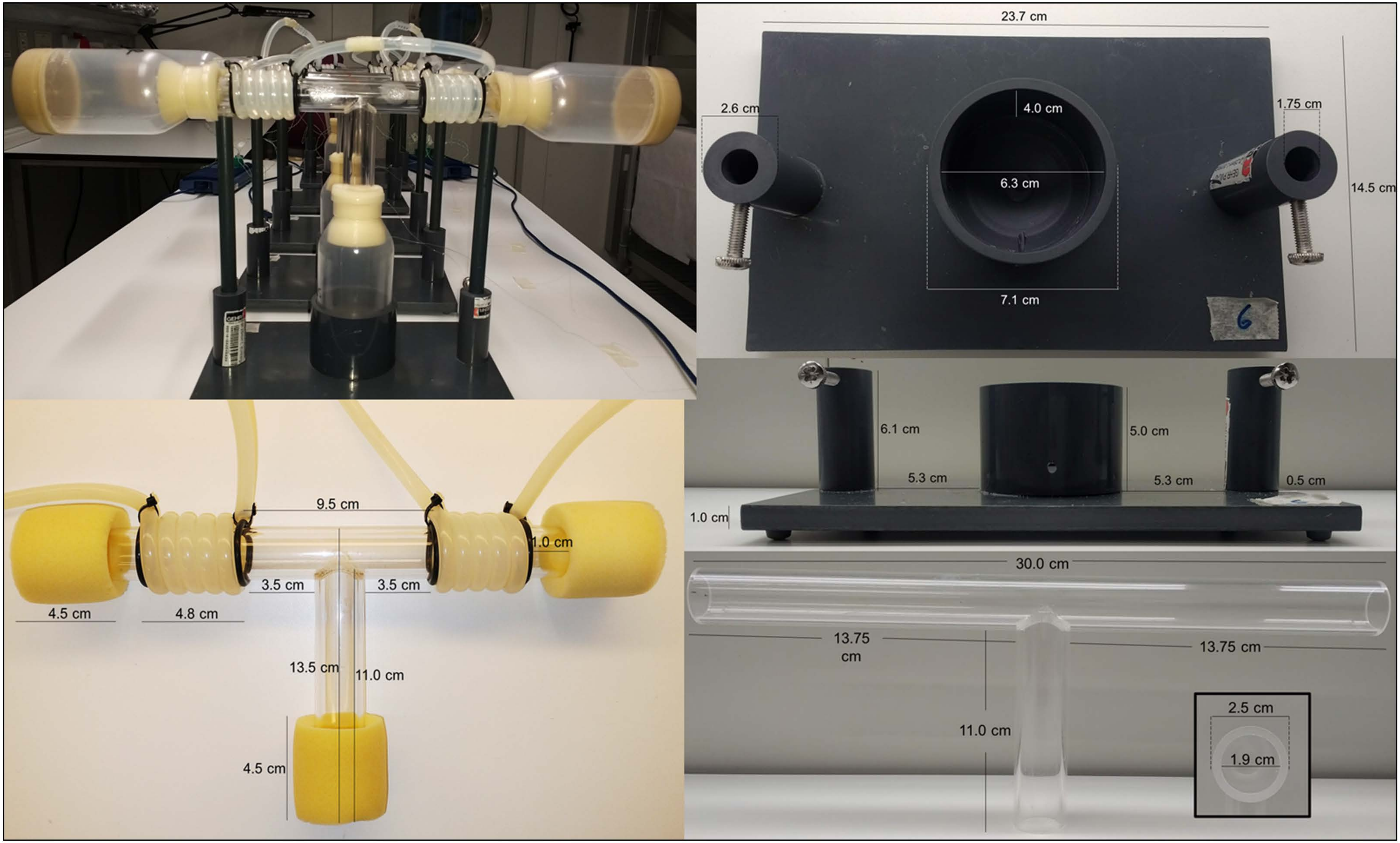
23.7 cm
2.6 cm
4.0 cm
1.75 cm
6.3 cm
14.5 cm
7.1 cm
6.1 cm
5.0 cm
5.3 cm
5.3 cm
0.5 cm
1.0 cm
9.5 cm
1.0 cm
3.5 cm
3.5 cm
4.5 cm
4.8 cm
13.5 cm
11.0 cm
4.5 cm
30.0 cm
13.75 cm
13.75 cm
11.0 cm
2.5 cm
1.9 cm

Figure S1

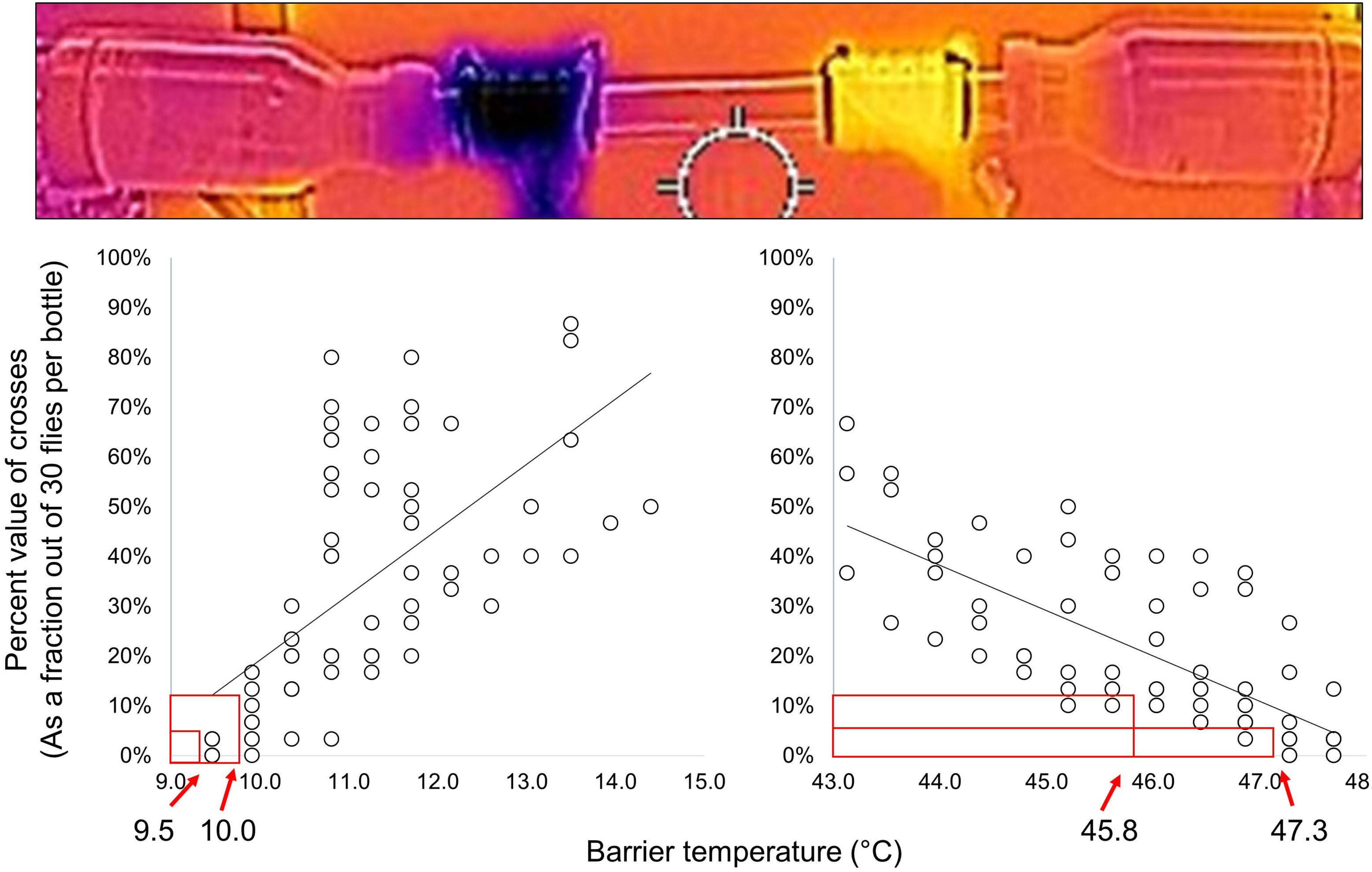
Percent value of crosses
(As a fraction out of 30 flies per bottle)
100%
90%
80%
70%
60%
50%
40%
30%
20%
10%
0%
9.0
10.0
11.0
12.0
13.0
14.0
15.0
43.0
44.0
45.0
46.0
47.0
48.0
9.5
10.0
45.8
47.3
Barrier temperature (°C)

Figure S2

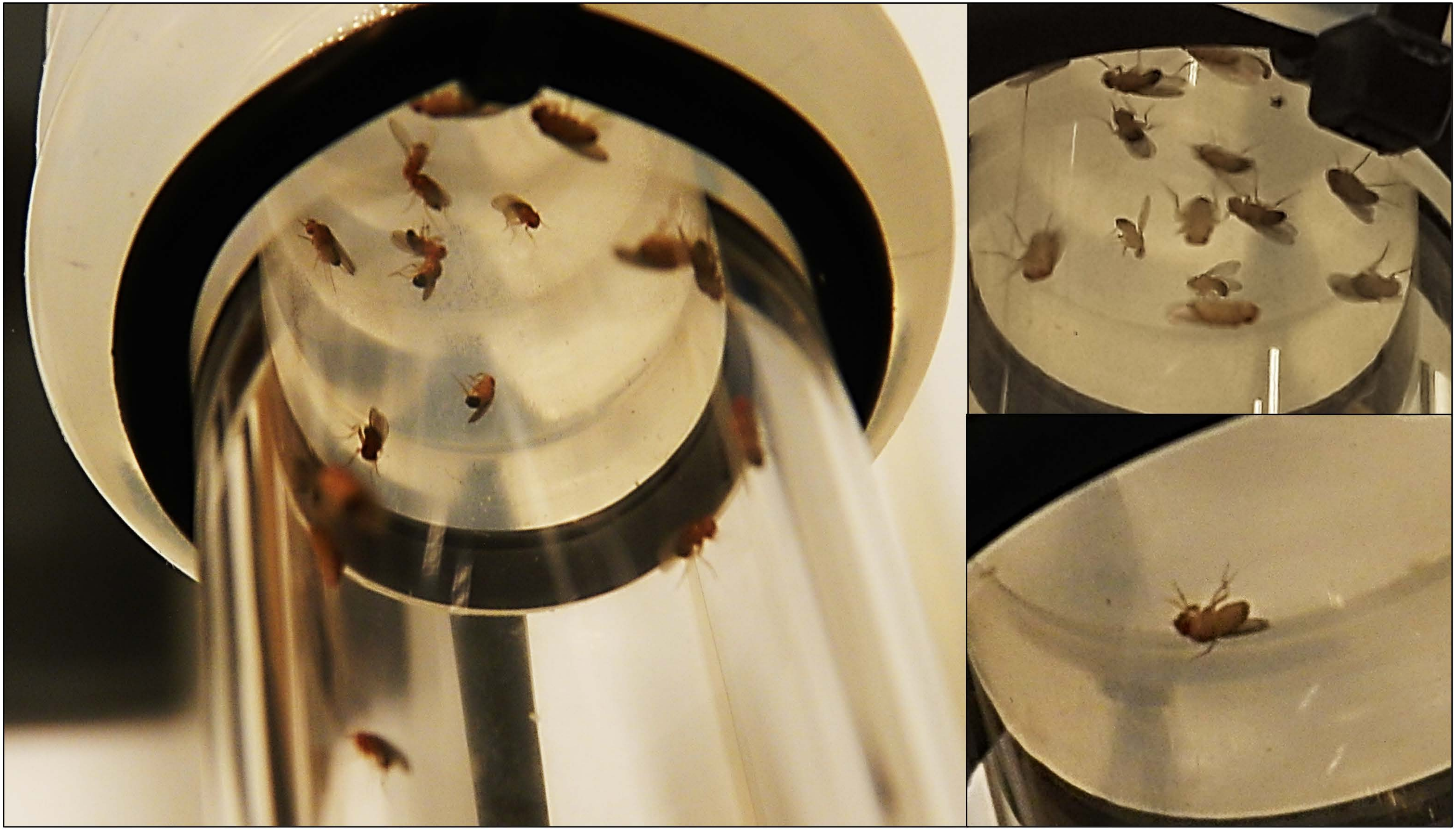

Figure S3

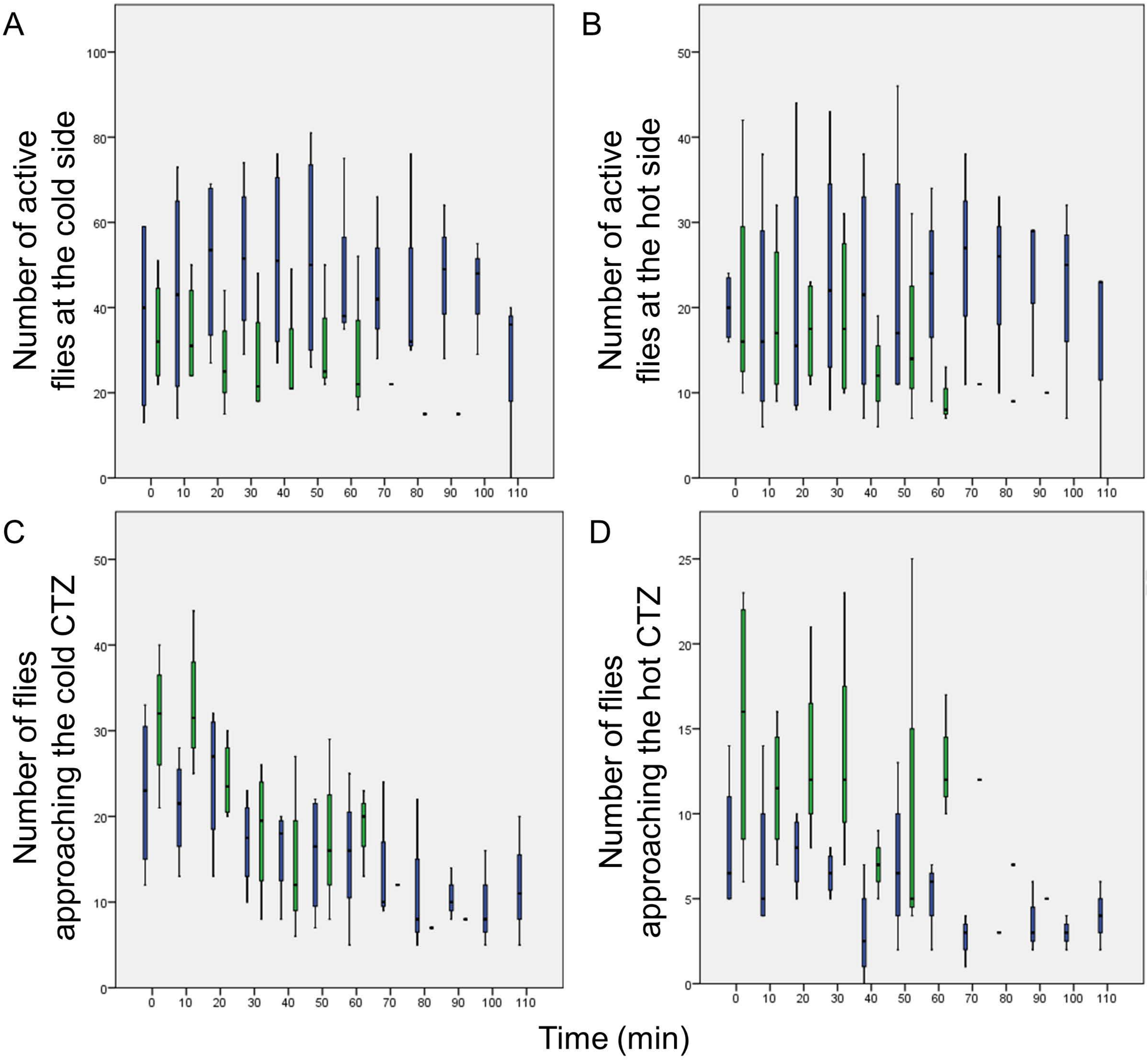
A
Number of active flies at the cold side
B
Number of active flies at the hot side
C
Number of flies approaching the cold CTZ
D
Number of flies approaching the hot CTZ
0 10 20 30 40 50 60 70 80 90 100 110
Time (min)

Figure S4

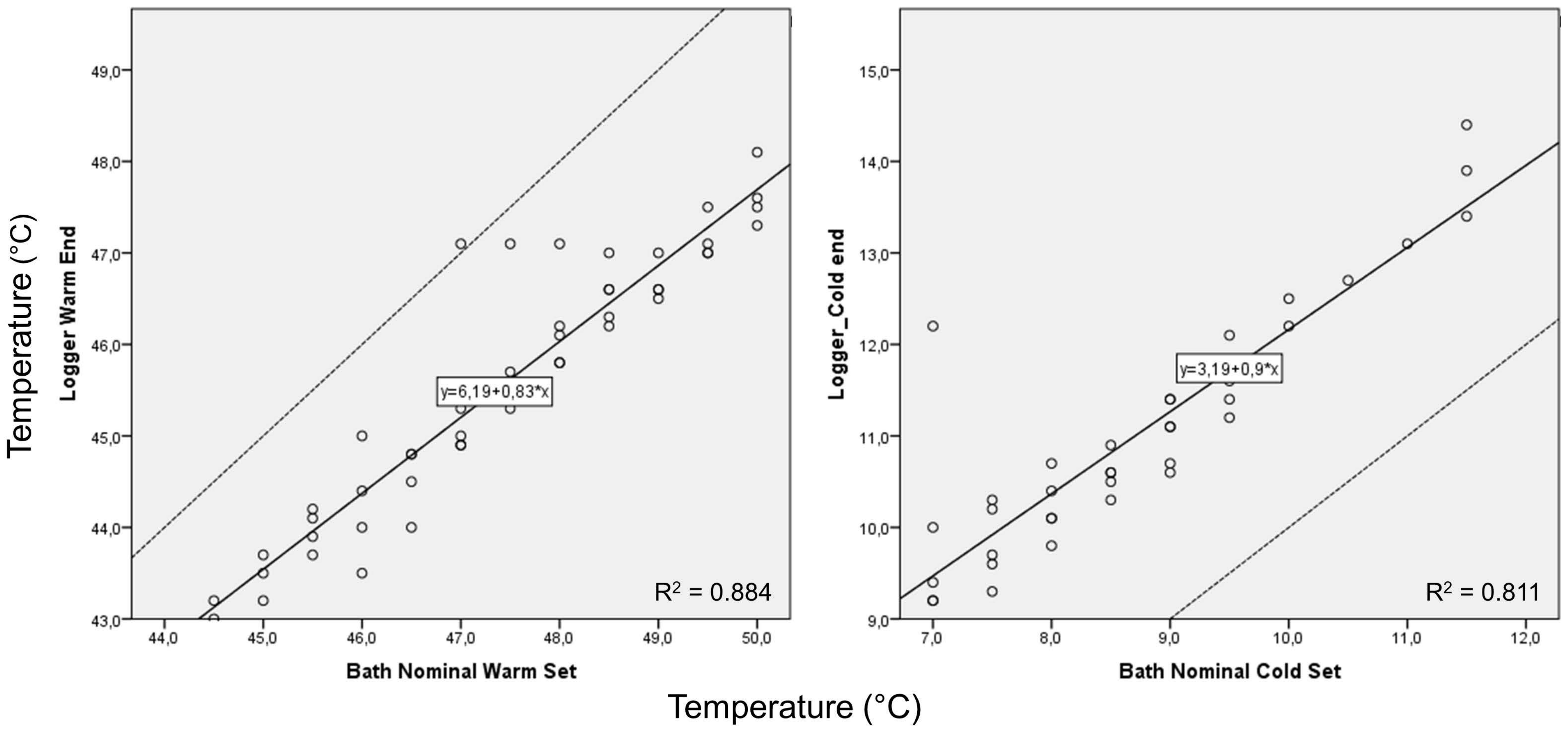
Temperature (°C)
Logger Warm End
49,0
48,0
47,0
46,0
45,0
44,0
43,0
44,0
45,0
46,0
47,0
48,0
49,0
50,0
Bath Nominal Warm Set
y=6,19+0,83*x
R² = 0.884
Logger_Cold end
15,0
14,0
13,0
12,0
11,0
10,0
9,0
7,0
8,0
9,0
10,0
11,0
12,0
Bath Nominal Cold Set
y=3,19+0,9*x
R² = 0.811
Temperature (°C)

Figure S5
